# Rapid Development of a Mucosal Nanoparticle Flu Vaccine by Genetic Engineering of Bacteriophage T4 using CRISPR-Cas

**DOI:** 10.1101/2022.06.13.495850

**Authors:** Mengling Li, Cen Chen, Xialin Wang, Pengju Guo, Helong Feng, Xueqi Zhang, Wanpo Zhang, Changqin Gu, Jingen Zhu, Guoyuan Wen, Venigalla B. Rao, Pan Tao

## Abstract

Mucosal vaccines that can induce local mucosal immune responses and combat the pathogens at entry sites are considered to be the most effective way to prevent infection. A universal platform that can be customized for development of mucosal vaccines against any given pathogen is therefore highly desired. Here, we demonstrate an efficient approach to develop nasal mucosal vaccines through genetic engineering of T4 phage to generate antigen-decorated nanoparticles. The antigen coding sequence was inserted into T4 genome in-frame at the C terminus of Soc (small outer capsid protein) using the CRISPR-Cas phage editing technology. During the propagation of recombinant T4 phages in *E. coli*, the Soc-antigen fusion proteins self-assemble on T4 capsids to form antigen-decorated nanoparticles that have intrinsic adjuvant activity and mucosal adhesive property. As a proof of concept, we showed that intranasal immunization with Flu viral M2e-decorated T4 nanoparticles efficiently induced local mucosal as well as systemic immune responses and provided complete protections against divergent influenza viruses in a mouse model. Potentially, our platform can be customized for any respiratory pathogen to rapidly generate mucosal vaccines against future emerging epidemics and pandemics.

## Introduction

The mucosal surfaces are entry sites for a majority of pathogens ^[1]^ and therefore represent ideal frontlines to combat infectious diseases. Vaccines that efficiently present antigens and elicit immune responses directly at local mucosal surfaces are highly preferred to prevent the invasion of pathogens at the first line of contact.^[2]^ However, most of the licensed vaccines are administrated through intramuscular (i.m.) immunization route, which mainly induce systemic immune responses and are inefficient in eliciting mucosal immunity.^[3, 4]^ While multiple routes such as oral, rectal, vaginal, and intranasal (i.n.) administration could be employed to induce mucosal immune responses, the intranasal delivery has been an attractive route due to the large surface area for absorption and the moderate local conditions.^[5, 6]^ However, all the limited nasal vaccines currently available for human and animal use are attenuated live viruses or bacteria, which might have safety concerns.^[7, 8]^

Subunit vaccines that use the immunogenic parts, particularly proteins, rather than the whole pathogen as vaccine antigens are safe alternatives, but it’s a challenge to develop intranasal subunit vaccines partially due to the lack of licensed mucosal adjuvants. Although a few adjuvants have been used in nasal vaccine clinical trials, recombinant subunit B of cholera heat-labile toxin is the only adjuvant licensed for mucosal vaccines.^[3]^ However, intranasal influenza vaccine (Nasalflu, Berna Biotech) used during 2000-2001 flu season in Switzerland was found associated with the increased risk of Bell’s palsy, which was likely due to the heat-labile toxin adjuvant included in the vaccine.^[9, 10]^ In addition, soluble protein antigens have short residence time after intranasal delivery due to the harsh conditions and mucociliary clearance.^[11, 12]^ The presence of thick layer of mucus further prevents the absorption of antigens and limits their recognition by antigen-presenting cells (APCs). Therefore, the vaccine platforms that can solve such limits for nasal delivery of antigens is desperately needed for mucosal vaccine development. To this end, a variety of nanoparticle vaccine platforms have been developed.^[13]^ However, adjuvant is still needed for the nanoparticles that lack intrinsic adjuvant activity.^[14]^

Recently, we developed a nanoparticle vaccine platform using T4 phage based on the high-affinity interactions between the small outer capsid protein (Soc) and capsid.^[15–17]^ Soc-fused antigens from plague, anthrax, influenza, and SARS-CoV2 were assembled on the surface of T4 capsid by incubating the recombinant fusions with *soc^-^* T4 phage *in vitro*.^[15, 16, 18, 19]^ The generated nanoparticles without any adjuvant induced robust systemic immune responses and provided complete protections against challenge in different animal models, indicating the potent intrinsic adjuvant activity of phage T4.^[15, 16]^

Recent studies indicated that phage T4 can be enriched in mucus layer on mucosal surfaces of human and all other tested animals through binding to mucin glycoproteins, which increases the chance to interact with their bacterial hosts.^[20]^ The mucoadhesive feature of phages along with their intrinsic adjuvant activities could be a defense mechanism of metazoan against invasive bacteria. After infection, phages use bacterial cell as a factory to manufacture and release hundreds of progeny phages, and an excess of progeny phages as well as bacterial lysis might recruit innate immune cells to clean off the bacteria. The mucoadhesive properties and intrinsic adjuvant activity of T4 phage led to our hypothesis that T4 phage could be an ideal mucosal vaccine platform, in addition to their ability to stimulate robust systemic immune responses against the delivered antigens.

Here, we report that phage T4 can be genetically engineered using CRISPR-Cas genome editing to rapidly develop a mucosal nanoparticle vaccine via assembly of antigens on T4 capsids *in vivo* (Figure 1). The coding sequence of extracellular domain of matrix protein 2 (M2e) of influenza A virus was inserted into T4 genome in-frame at the C terminus of Soc gene, which was used as an anchor for *in vivo* display (Figure S1).^[21]^ During the propagation of recombinant phage T4 in *E. coli*, expressed Soc-M2e proteins efficiently self-assembled on T4 capsids to form M2e-T4 nanoparticles (Figure 1A). Significantly, i.n. administration of M2e-T4 nanoparticles elicited robust mucosal as well as systemic immune responses without any exogenous adjuvant and conferred mice enhanced protection against divergent influenza virus challenges when compared to the i.m. vaccination. Mechanistic studies indicated that T4 nanoparticles remained in the respiratory tract for at least 26 days after i.n. administration, which significantly enhanced antigen uptake by pulmonary antigen-presenting cells (APCs) and promote adaptive immune responses (Figure 1B). High levels of secretory IgA and CD4^+^ T cell immune responses, including effector memory T cells (T_EM_) and tissue-resident memory T cells (T_RM_), against M2e were induced in the respiratory mucosa, which mediated cross-protective mucosal immunity against divergent influenza viruses.^[22, 23]^ Our findings demonstrate that T4 nanoparticle is a promising mucosal delivery platform that can be used to rapidly develop mucosal vaccines against the emerging respiratory pathogens.

**Figure 1.**
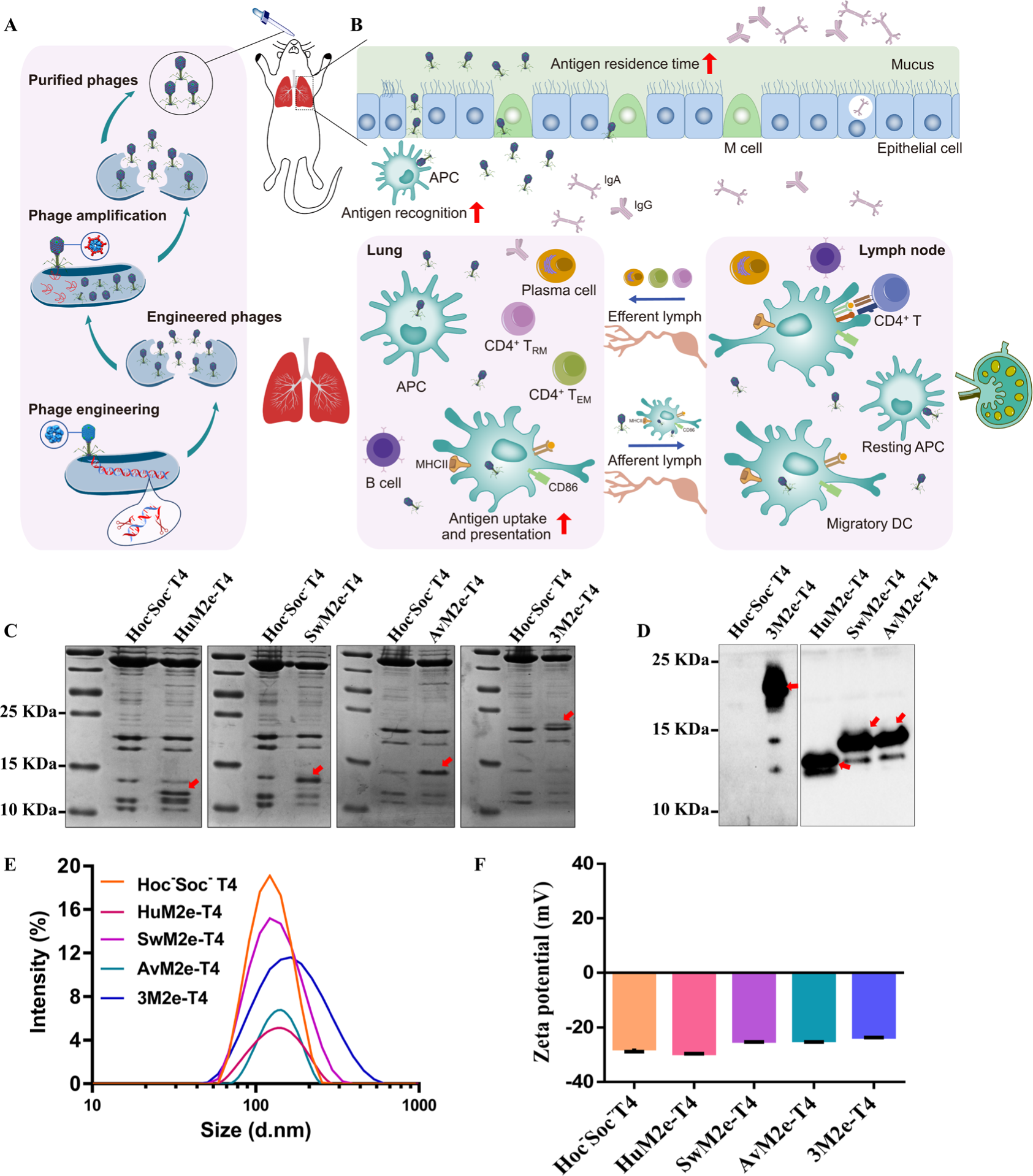
A mucosal vaccine platform based on T4 phage nanoparticle. Schematic showing the generation of recombinant T4 phages and preparation of antigen-decorated T4 nanoparticles *in vivo*. **(A)** that can persist in mucus layer and lead to the prolonged exposure of the antigen to immune system to enhance mucosal immune responses **(B)**. The enlarged views show the single capsomer of phages *hoc^-^soc^-^* T4 (**A, bottom**) or antigen-decorated T4 (**A, middle**). **(C-F)** Generation and characterization of M2e-T4 nanoparticles. The presence of M2e antigens on T4 nanoparticles was determined by SDS-PAGE **(C)** and Western blot **(D)** analysis of recombinant phages, HuM2e-T4, SwM2e-T4, AvM2e-T4, and 3M2e-T4. The *hoc^-^soc^-^* T4 phage was used as a control. Red arrows indicate Soc-HuM2e, Soc-SwM2e, Soc-AvM2e, and Soc-3M2e. Particle size distribution **(E)** and zeta potential **(F)** of *hoc^-^soc^-^* T4 (orange line), HuM2e-T4 (red line), SwM2e-T4 (purple line), AvM2e-T4 (green line), and 3M2e-T4 nanoparticles (blue line) were determined as described in the Materials and Methods.

## 2. Results

### 2.1 Assembly of 3M2e decorated T4 nanoparticles *in vivo* using genetically engineered phage T4

Previously, we used an *in vitro* assembly system to display antigen-Soc fusion proteins on T4 capsid by mixing the fusion proteins with *Hoc^-^Soc^-^* T4 phages.^[15–17, 24]^ The highly efficient phage genome editing technology recently developed using CRISPR-Cas allowed us to display foreign proteins on T4 capsid *in vivo* by inserting the coding sequence into T4 genome in-frame at the C terminus of Soc gene.^[19, 21, 25]^ During the propagation of recombinant T4 phages in *E. coli*, expressed Soc-antigen proteins are expected to assemble on T4 capsids to form antigen-decorated nanoparticles. Assembly of such nanoparticle *in vivo* will avoid the purification of Soc-antigen proteins and the subsequent *in vitro* assembly, greatly accelerating the development of nanoparticle vaccines. To assemble M2e proteins on T4 nanoparticles *in vivo*, the coding sequences of M2e of human, swine, and avian influenza viruses were inserted into T4 genome individually to generate three recombinant T4 phages, HuM2e-T4, SwM2e-T4, and AvM2e-T4 (Figure S1). The 3M2e gene containing three tandem copies of M2e from human, swine, and avian influenza viruses was also inserted into T4 genome (3M2e-T4) to cover divergent influenza viruses (Figure S1).

Sodium dodecyl sulfate-polyacrylamide gel electrophoresis (SDS-PAGE) analysis of the CsCl-purified recombinant T4 phages showed the presence of specific bands with a molecular mass between 10 kDa and 25 kDa in recombinant phages (Figure 1C, red arrows). These bands corresponding to Soc-HuM2e, Soc-SwM2e, Soc-AvM2e, and Soc-3M2e were further confirmed by Western blot using anti-M2e antibodies (Figure 1D). The average diameter of M2e decorated T4 nanoparticles is about 127.7-148.6 nm, which is higher than that of *hoc^-^soc^-^*T4 phage (118.6 nm) (Figure 1E). The Zeta-potential of M2e decorated T4 nanoparticles was around -30 to -23 mV, which is comparable to that of *hoc^-^soc^-^* T4 phages (−27.7±1.11 mV), indicating the negatively charged surface of T4 nanoparticles in suspension (Figure 1F). To evaluate the stability of the M2e-T4 nanoparticles, we analyzed the CsCl-purified T4 phages, which were stored at 4℃ for ten months, by SDS-PAGE. The Soc-M2e and Soc-3M2e bands can be observed on SDS-PAGE gel, indicating the stability of M2e decorated T4 nanoparticles (Figure S2).

### 2.2 I.m. immunization of mice with 3M2e-T4 nanoparticles induced complete protection against homologous influenza virus but partial protection against heterologous virus challenges

We have shown previously that 3M2e-T4 nanoparticles assembled *in vitro* provided complete protection to mice against homologous influenza virus challenge when administrated through i.m. route.^[16]^ To determine the protective efficacy of the nanoparticles prepared above using *in vivo* assembly, mice were intramuscularly immunized three times with 3M2e-T4 or a mixture of HuM2e-T4, SwM2e-T4, and AvM2e-T4 on day 0, 14, and 28 (Figure 2A). Mice immunized with a similar amount of phage T4 or Soc-3M2e soluble proteins were used as controls. Sera were collected as indicated in Figure 2A, and M2e-specific antibodies were determined using enzyme-linked immunosorbent assay (ELISA). As shown in Figure 2B, the mixture of HuM2e-T4, SwM2e-T4, and AvM2e-T4 induced a significantly higher level of M2e-specific IgG than soluble Soc-3M2e proteins, while the 3M2e-T4 nanoparticles elicited the highest titers with the end point titer of ∼2×10^5^. As expected, the antibody levels in the phage T4 control group were negligible. The soluble Soc-3M2e proteins mainly induced IgG1 antibodies, while the 3M2e-T4 nanoparticles and the mixture of HuM2e-T4, SwM2e-T4, and AvM2e-T4 induced comparable levels of M2e-specific IgG1 and IgG2a (Figure 2C, D), indicating that T4 nanoparticles induced balanced Th1/Th2 immune responses. Interestingly, only the 3M2e-T4 nanoparticles induced M2e-specific IgA (Figure 2E).

**Figure 2.**
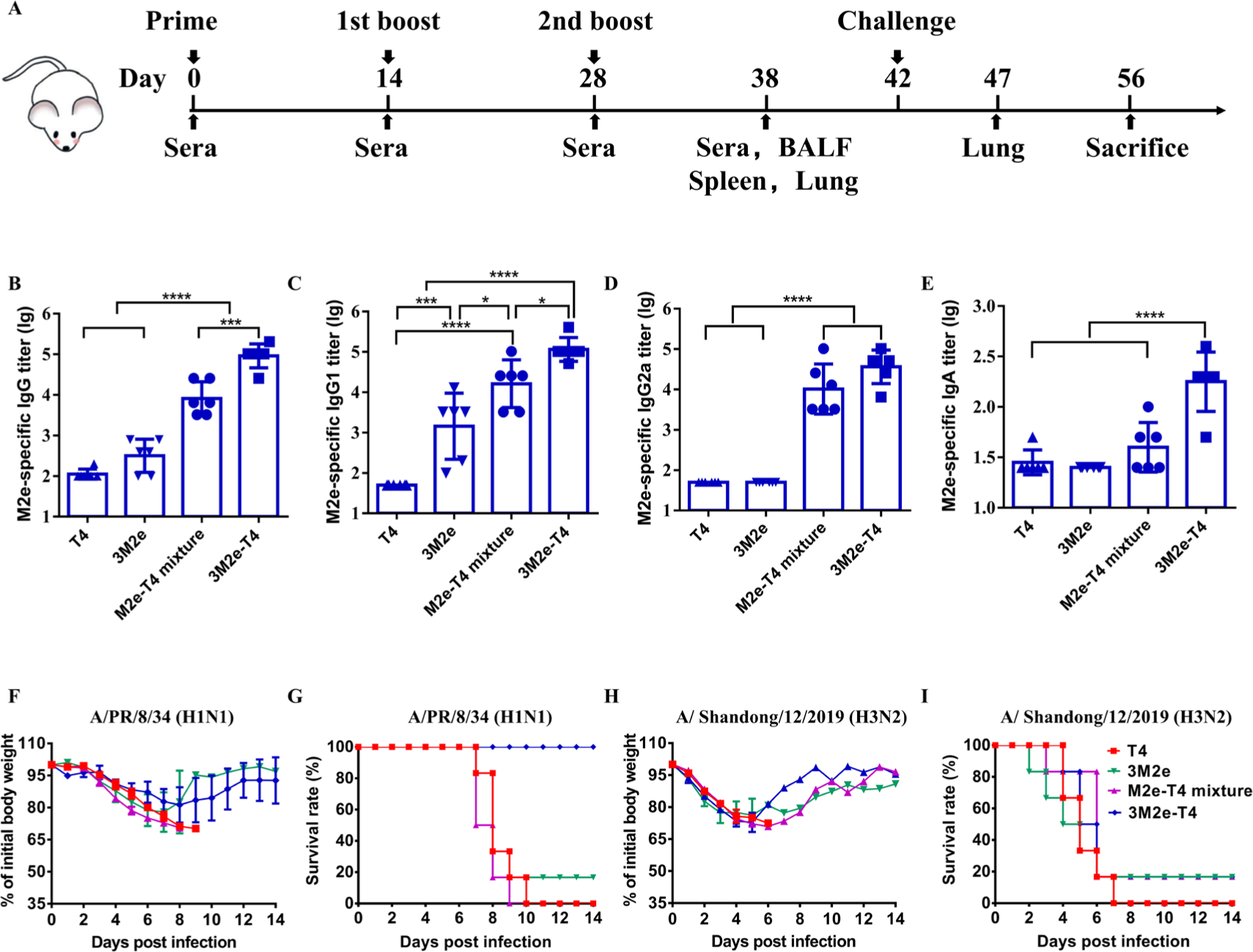
Immunogenicity and protective efficiency of *in vivo* assembled 3M2e-T4 nanoparticles via i.m administration. Mice were immunized three times and sera were collected according to the scheme **(A).** The titers of total IgG **(B)**, IgG1 **(C)**, IgG2a **(D)**, and IgA **(E)** antibodies against M2e peptides were measured by ELISA. The protective efficacy was determined by challenge with 5×LD_50_ of homologous influenza virus A/Puerto Rico/8/1934 (**F, G**) and 3×LD_50_ of heterologous influenza virus A/Shandong/12/2019 **(H, I)**. Weight loss **(F, H)** and survivals **(G, I)** of immunized mice (n=6) were monitored daily for 14 days. Data represent mean ± S.D. *P < 0.05; ***P < 0.001; ****P < 0.0001.

The efficacy of T4 nanoparticles was tested by challenge of immunized mice two weeks after last vaccination with 5LD_50_ of A/Puerto Rico/8/34 (H1N1) (A/PR/8/34) and 3LD_50_ of A/ duck/Shandong/12/2019 (H3N2) respectively. The A/PR/8/34 was used as a homologous virus while the latter was used as a heterologous virus, based on the sequence identity of M2e genes inserted into T4 genome (Figure S1D). All mice experienced weight loss but the mice immunized with 3M2e-T4 nanoparticles regained their body weight 9 days post challenge (Figure 2F). All the T4 control mice and 5 of 6 mice vaccinated with Soc-3M2e soluble proteins died within 10 days after challenge, whereas the 3M2e-T4 nanoparticles provided complete protection (Figure 2G). Unexpectedly, all mice immunized with the mixture of HuM2e-T4, SwM2e-T4, and AvM2e-T4 died 9 days after the challenge (Figure 2G). This might because the structural character of Soc protein on T4 capsid, which sticks to the capsid surface as a “glue” between neighboring hexameric capsomers (Figure 1A),^[26]^ and the short peptide (M2e) fused to Soc was not well exposed to immune system compared to the long peptide (3M2e). The heterologous immune protection of T4 nanoparticles was evaluated by challenge of immunized mice with 3LD_50_ of A/duck/Shandong/12/2019 (H3N2). All T4 control mice died 7 days post challenge whereas 16.7% of mice immunized with Soc-3M2e soluble proteins or antigen-decorated T4 nanoparticles were survival (Figure 2H-I). We focused on the 3M2e-T4 nanoparticle in our following studies because of its higher protective efficacy against homologous challenge compared with HuM2e-T4, SwM2e-T4, and AvM2e-T4.

### 2.3 I.n. immunization of mice with 3M2e-T4 nanoparticles elicited complete protection against homologous and heterologous influenza virus challenges

Since influenza A viruses mainly infect respiratory tract, the local mucosal immune responses in the respiratory tract elicited by intranasal vaccines are optimally positioned and could provide better protection against infections. To determine whether T4 phage can be used as an intranasal vaccine platform for antigen delivery, mice were immunized intranasally three times with 3M2e-T4 nanoparticles as shown in Fig 2A. PBS, T4 phages, and the soluble Soc-3M2e were used as controls. When challenged with 5LD_50_ of A/PR/8/34, all mice immunized with 3M2e-T4 nanoparticles through i.n. route survived and showed less weight loss compared to the mice immunized via i.m. route (Figure 3A-B, Figure 2F-G). In contrast, all mice in the control groups showed continuous weight loss and died within 10 days post challenge (Figure 3A-B). The protection effect of 3M2e-T4 was also supported in a separated experiment by pathological observation of the lungs of vaccinated mice, which showed relatively normal alveolar wall thickness and negligible inflammatory infiltration 5 days after challenge (Figure 3E). Instead, the alveolar walls of mice in the control groups were severely thickened (Figure 3E, row II), and the bronchi (Figure 3E, row III) and pulmonary blood vessels (Figure 3E, row IV) were infiltrated by a large number of inflammatory cells.

**Figure 3.**
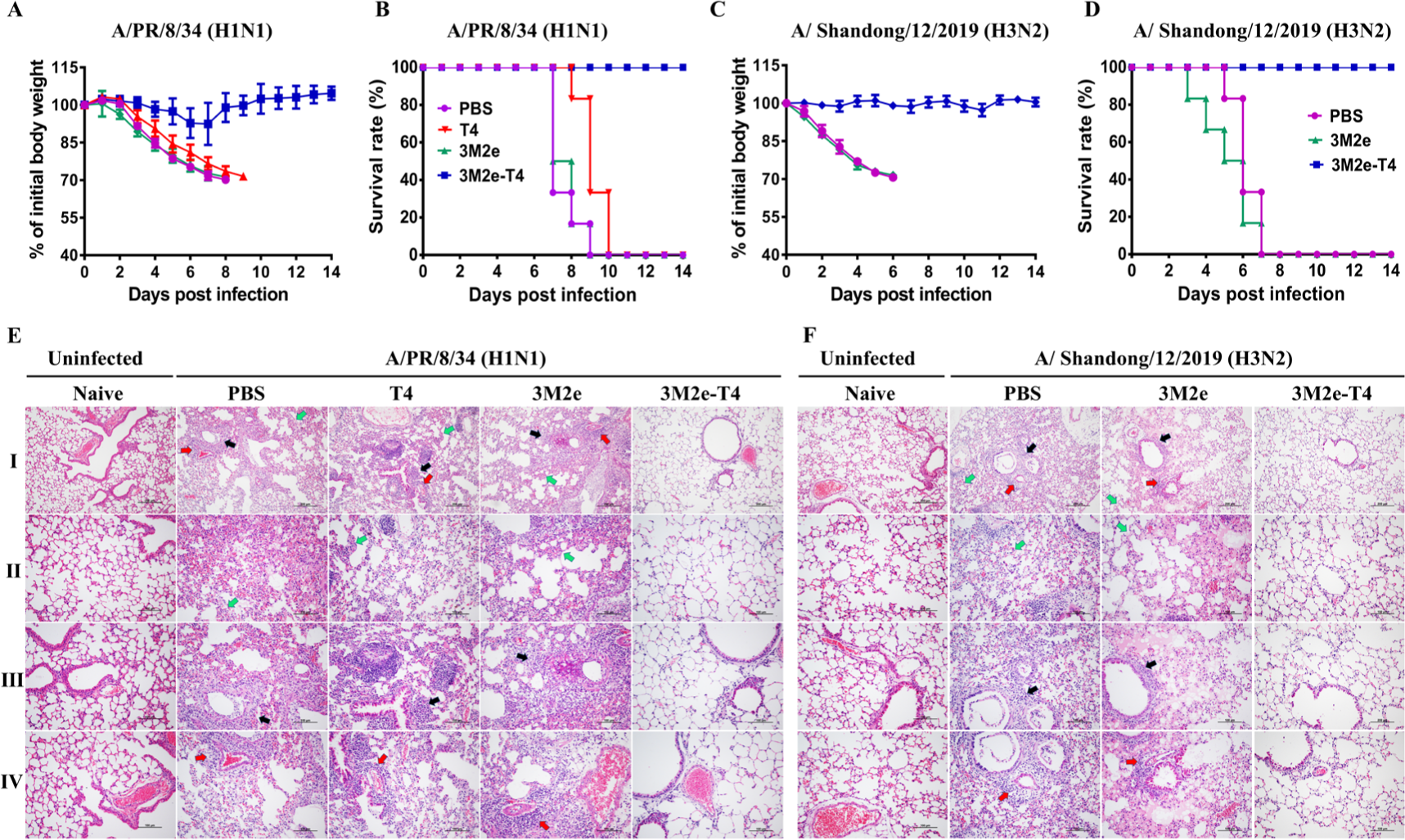
I.n. immunization with 3M2e-T4 nanoparticles conferred mice complete protection against homologous and heterologous influenza virus challenges. Experimental scheme of immunization and challenge was shown in Figure 2A. Weight loss **(A, C)** and survivals **(B, D)** of immunized mice (n=6) were monitored daily for 14 days after challenge with 5×LD_50_ of homologous influenza virus A/PR/8/34 **(A, B)** and 3×LD_50_ of heterologous influenza virus A/Shandong/12/2019 **(C, D)**. Pathological analysis of lung from immunized mice (n=3) at 5 days after challenge with A/PR/8/34 **(E)** and A/Shandong/12/2019 **(F)**. The representative lung tissue sections were shown (column Ⅰ, scale bar=200 μm; columns Ⅱ-Ⅳ show the zoom-ins of column I, scale bar=100 μm). The main pathological findings are the thickening of alveolar wall (column Ⅱ, green arrows) and infiltration of inflammatory cells around the bronchi (column Ⅲ, black arrows) and pulmonary vessels (column Ⅳ, red arrows).

In contrast to i.m. immunization, i.n. delivery of 3M2e-T4 nanoparticles induced complete protection against challenge with 3LD_50_ of heterologous influenza virus A/Shandong/12/2019 (H3N2), and no significant body weight loss was observed (Figure 3C-D). As expected, all the control mice suffered from severe weight loss and died within 7 days post challenge. Similarly, the protection of 3M2e-T4 nanoparticles against heterologous challenge was further evaluated in a separated experiment by pathological analysis of the lungs of the immunized mice 5 days post-challenge (Figure 3F). Mice in the control groups exhibited massive lung lesions and obvious infiltration of inflammatory cells around the bronchi (Figure 3F, row Ⅲ) and pulmonary blood vessels (Figure 3F, row Ⅳ). The alveolar walls were markedly thickened due to the inflammation. However, no significant lung lesions were observed in mice immunized with 3M2e-T4 nanoparticles (Figure 3F). Taken together, these results demonstrated that i.n. immunization with 3M2e-T4 nanoparticles conferred mice enhanced protections against both homologous and heterologous viruses challenge, indicating T4 phage could be a good intranasal vaccine platform to deliver antigens.

### 2.4 I.n. immunization with 3M2e-T4 nanoparticles elicited similar systemic immune responses compared to intramuscular delivery

I.n. vaccination can induce local mucosal as well as systemic immune responses.^[27]^ To determine why i.n. immunization increases the protective efficacy of 3M2e-T4 nanoparticles compared to the i.m. administration, we first analyzed systemic humoral immune responses induced by two different immunization routes. The 3M2e-T4 elicited robust M2e-specific IgG in sera after i.n. immunization, with the endpoint titer of ∼1×10^5^ (Figure 4A). As expected, the titers of M2e-specific IgG in PBS and phage T4 groups were negligible while the soluble Soc-3M2e induced a quite low level of IgG (Figure 4A). As observed in i.m. immunization, i.n. vaccination of 3M2e-T4 also induced balanced anti-M2e IgG1 and IgG2a (Figure 4B and C). However, there were no significant differences in total IgG, IgG1, and IgG2a between i.m. and i.n. immunizations (Figure 4A-C). The M2e-specific IgA antibodies in sera were also induced by 3M2e-T4 nanoparticles when administrated via i.n. route, and no significant difference was observed between i.m. and i.n. routes (Figure 4D). Since anti-M2e antibodies mediated immune protection mainly depends on antibody-dependent cellular cytotoxicity (ADCC) and antibody-dependent cellular phagocytosis (ADCP),^[28]^ we determined the ability of M2e-specific IgG to recognize Madin-Darby canine kidney (MDCK) cells infected by influenza viruses. The results revealed that M2e-specific IgG antibodies elicited via i.m. and i.n. routes can similarly recognize MDCK cells infected with A/PR/8/34 and A/duck/Shandong/12/2019 (H3N2) respectively (Figure 4E). These results indicated that the enhanced protection of 3M2e-T4 nanoparticles via i.n. route was not because of the systemic humoral immune responses.

**Figure 4.**
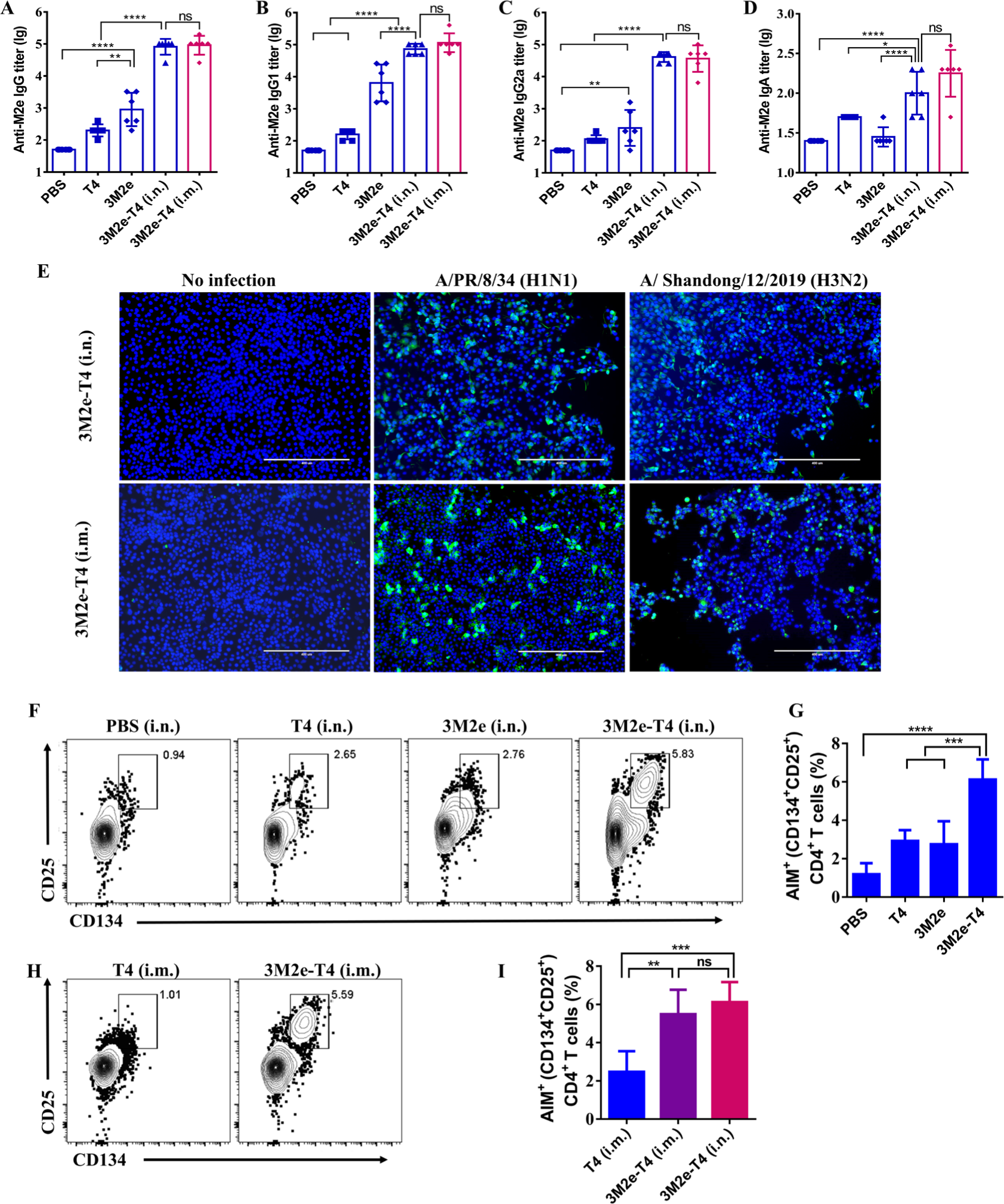
I.n. and i.m. immunization of 3M2e-T4 nanoparticles elicited similar systemic immune responses. Scheme of immunization was shown in Figure 2A, and sera were collected at 10 days after the last immunization. M2e-specific IgG **(A)**, IgG1 **(B)**, IgG2a **(C)**, and IgA **(D)** antibodies titers were determined by ELISA assay. **(E)** The binding of M2e-specific IgG to influenza virus-infected MDCK cells was determined by indirect immunofluorescence assay. Cells were infected with A/PR/8/34 or A/Shandong/12/2019 at MOI of 1. M2e-specific CD4^+^ T cells (**F-I**) were analyzed by AIM assays as described in the Materials and Methods. The spleen lymphocytes were isolated at 10 days after the third immunization, and M2e-specific CD4+ T cells were analyzed using flow cytometry after stimulation with M2e peptides. Representative plots of M2e-specific CD4^+^ T cells of immunized mice via i.n. **(F)** and i.m. route **(H)**. Data were represented as mean ± S.D. *P < 0.05; **P < 0.01; ***P < 0.001; ****P < 0.0001.

To determine systemic cellular immune responses, mice were vaccinated as shown in Fig 2A with 3M2e-T4 nanoparticles via i.m. and i.n. routes respectively, and spleen lymphocytes were isolated 10 days after the last immunization. Since M2e does not contain a CD8^+^ T cell epitope, we focused on CD4^+^ T cells due to their critical roles in M2e-induced immune protection.^[29]^ M2e-specific CD4^+^ T cells were analyzed by T cell receptor (TCR) dependent activation-induced marker (AIM) assays, which have been widely used to identify antigen-specific CD4^+^ T cells.^[30–34]^ Unlike intracellular cytokine staining and ELISPOT assays, which only determine the antigen-specific T cells that secrete specific cytokines, AIM assay can simultaneously identify the total antigen-specific T cells.^[34, 35]^ The spleen lymphocytes were stimulated with M2e peptide and analyzed by flow cytometry after staining with fluorescent antibodies against AIM markers, CD25 and CD134.^[32–34]^ As shown in Figure 4F and G, i.n. immunization with 3M2e-T4 nanoparticles induced a significantly higher level of M2e-specific CD4^+^ T cells (CD25^+^CD134^+^) immune responses compared to that of the soluble Soc-3M2e protein (p<0.001, ANOVA). However, there is no significant difference between i.m. and i.n. immunization routes (Figure 4H, I). As expected, mice in PBS and T4 phage control groups generated only background level of M2e-specific CD4^+^ T cell responses (Figure 4F-I). Taken together, these results indicated that enhanced protection of 3M2e-T4 through i.n. delivery cannot be explained by systemic humoral and cellular immune responses.

### 2.5 Intranasal immunization with 3M2e-T4 nanoparticles induced robust humoral and cellular responses at local mucosal site

The local immune responses at the pathogen encounter sites have many advantages to fight against infections,^[22]^ and mucosal immunization of antigens is necessary for the induction of local immune responses.^[36, 37]^ To determine whether i.n. delivery of 3M2e-T4 nanoparticles can efficiently induce local immune responses in addition to systemic immune responses, we first determined the secretory IgA (sIgA) in the lungs, which plays critical roles in mucosal defense.^[37, 38]^ The bronchoalveolar lavage fluid (BALF) was obtained 10 days after the last i.n. vaccination, and M2e-specific IgA as well as IgG were determined. Mice immunized with 3M2e-T4 nanoparticles via i.m. route was used as a comparison group. As shown in Figure 5A, i.n. vaccination with 3M2e-T4 nanoparticles induced a significantly higher level of M2e-specific IgA than that in groups PBS, soluble Soc-3M2e protein, and phage T4. Similarly, i.n. vaccination with 3M2e-T4 also efficiently elicited M2e-specific IgG antibodies compared to the control groups (Figure 5B). However, in contrast to the i.n. administration, i.m. immunization with 3M2e-T4 nanoparticles induced comparable levels of M2e-specific IgG in BALF as that of i.n. immunizations (Figure 5C) but failed to induce M2e-specific IgA (Figure 5D), which is consistent with our previous study.^[16]^

**Figure 5.**
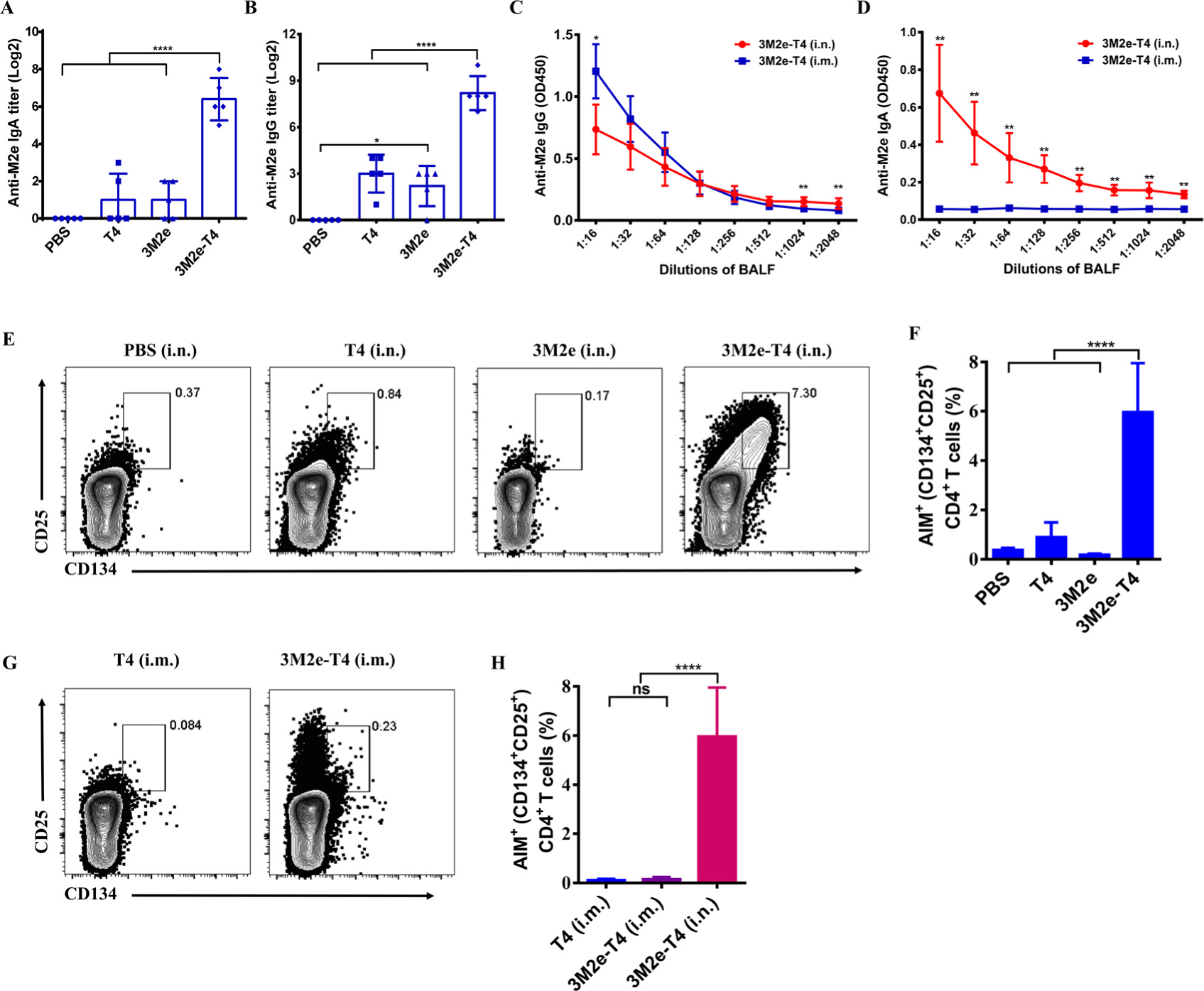
I.n immunization of 3M2e-T4 nanoparticles elicited strong mucosal humoral and CD4^+^ T cell immune responses in lung. Bronchoalveolar lavage fluid (BALF) was collected at 10 days after third immunization. M2e-specific IgA **(A)** and IgG **(B)** antibodies titers were measured by ELLSA assay. The differences in M2e-specific IgG **(C)** and IgA **(D)** antibodies between i.n. and i.m. immunization were shown. Lung M2e-specific CD4^+^ T cells of intranasally **(E, F)** and intramuscularly **(G, H)** immunized mice were measured by AIM assays, and representative flow cytometry plots were shown (**E, G**). Data were represented as mean ± S.D. *P < 0.05; **P < 0.01; ****P < 0.0001.

To determine CD4^+^ T cell responses in lungs, pulmonary lymphocytes were isolated 10 days after last immunization, and the M2e-specific CD4^+^ T cells were determined using AIM assay as described above. Mice intranasally immunized with 3M2e-T4 nanoparticles accumulated significantly higher levels of M2e-specific CD4^+^ T cells in the lungs compared to the mice immunized with PBS, soluble Soc-3M2e protein or T4 phages (Figure 5E, F) (p<0.0001, ANOVA). Interestingly, we found i.m. delivery of 3M2e-T4 nanoparticles showed much less accumulation of M2e-specific CD4^+^ T cells in the local lungs compared to i.n. administration (Figure 5G, H) (p<0.0001, ANOVA). These results also indicated that i.n. but not i.m. delivery of 3M2e-T4 nanoparticles can induce robust CD4^+^ T cell responses in local lung, which is consistent with previous findings that mucosal immunization of antigens is necessary for the induction of local T cell responses.^[36]^

Memory CD4^+^ T cells are central regulators of the adaptive immune responses and are critical for the long-term protection against influenza virus infection.^[22, 29]^ To further analyze M2e-specific CD4^+^ T cells in lungs, CD4^+^ T cells were pre-gated on CD44 and CD62L to identify effector memory cells (T_EM_, CD62L^low^CD44^hi^) and central memory cells (T_CM_, CD62L^hi^CD44 ^hi^).^[39]^ The former prefers homing non-lymphoid peripheral tissues such as lung during circulation, while T_CM_ cells have a similar circulation path as naive CD4^+^ T cells. As shown in Figure 6A-B, significantly higher levels of M2e-specific T_EM_ cells were observed in lungs of mice intranasally immunized with 3M2e-T4 nanoparticles compared with that of PBS, T4 phage, and soluble Soc-3M2e protein groups. Significantly, i.m. vaccination of 3M2e-T4 nanoparticles failed to accumulate M2e-specific T_EM_ cells in local lungs (Figure 6D, E). Similarly, i.n. immunization with 3M2e-T4 nanoparticles also induced more M2e-specific T_CM_ cells than i.m. administration did (Figure 6A, C, D, F).

**Figure 6.**
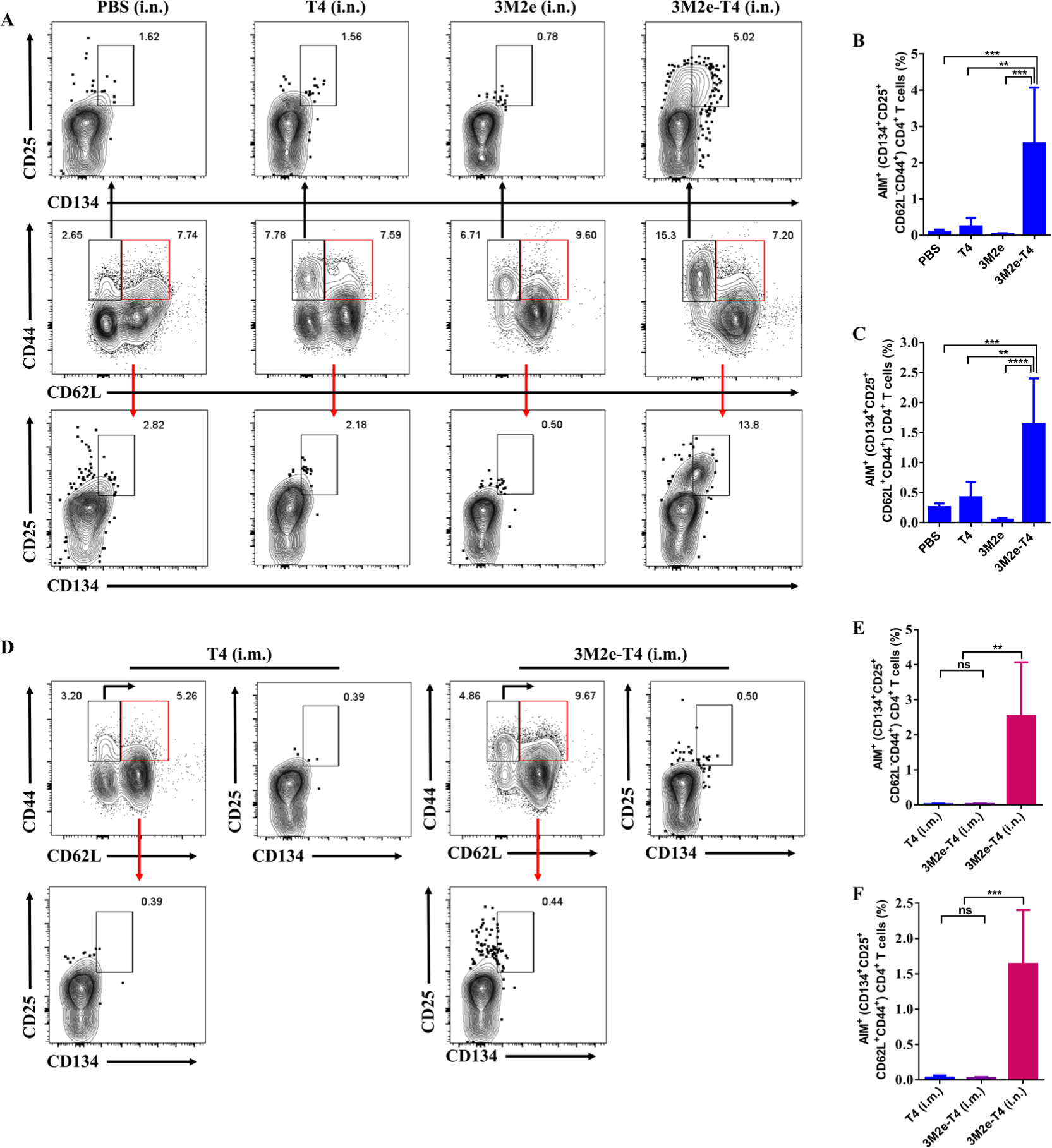
I.n. immunization of 3M2e-T4 nanoparticles elicited M2e-specific CD4^+^ T_EM_ and CD4^+^ T_CM_ in lungs. The frequencies (%) of CD4^+^ T_EM_ and T_CM_ in lungs were determined at 10 days after administration via i.n. **(A-C)** or i.m. **(D-F)** route. Representative flow cytometry plots showed i.n. immunization with 3M2e-T4 nanoparticles elicited higher M2e-specific CD4^+^ T_EM_ **(A, top row**) and T_CM_ (**A, bottom row**) than control groups that immunized intranasally with PBS, T4, and soluble 3M2e protein respectively. However, i.m. immunization with 3M2e-T4 nanoparticles failed to elicit M2e-specific CD4^+^ T_EM_ (**D**, top row) and T_CM_ (**D**, bottom row). The differences in M2e-specific CD4^+^ T_EM_ (**E**) and T_CM_ (**F**) cells between i.n. and i.m. administrations were represented as mean ± S.D. ** P<0.01; ***P < 0.001; ****P< 0.0001.

The third subset of memory CD4^+^ T cells, tissue-resident memory T cells (T_RM_), which persisted in the lung long-term after influenza virus infection, are believed to be critical defenders in the mucosal tissues upon virus re-encounter.^[22]^ To determine the M2e-specific T_RM_ cells, the total lung CD4^+^ T cells were pre-gated on CD69 and CD11a, which have been widely accepted as the common markers of T_RM_ (CD69^+^CD11a^+^).^[40, 41]^ Results revealed that mice immunized with 3M2e-T4 nanoparticles generated significantly higher levels of M2e-specific T_RM_ cells than mice immunized with the PBS, soluble Soc-3M2e protein or T4 phages (Figure 7A, B). Most importantly, significant difference was found between i.n. and i.m. delivery (p<0.01, ANOVA), the latter inducing only background levels of M2e-specific T_RM_ cells in the lungs (Figure 7C, D). Taken together, the above sets of data strongly indicate that mucosal anti-M2e IgA and local lung M2e-specific CD4^+^ T cells, including T_EM_ and T_RM_ cells, could be the critical reasons for the enhanced protection of 3M2e-T4 via i.n. immunization route.

**Figure 7.**
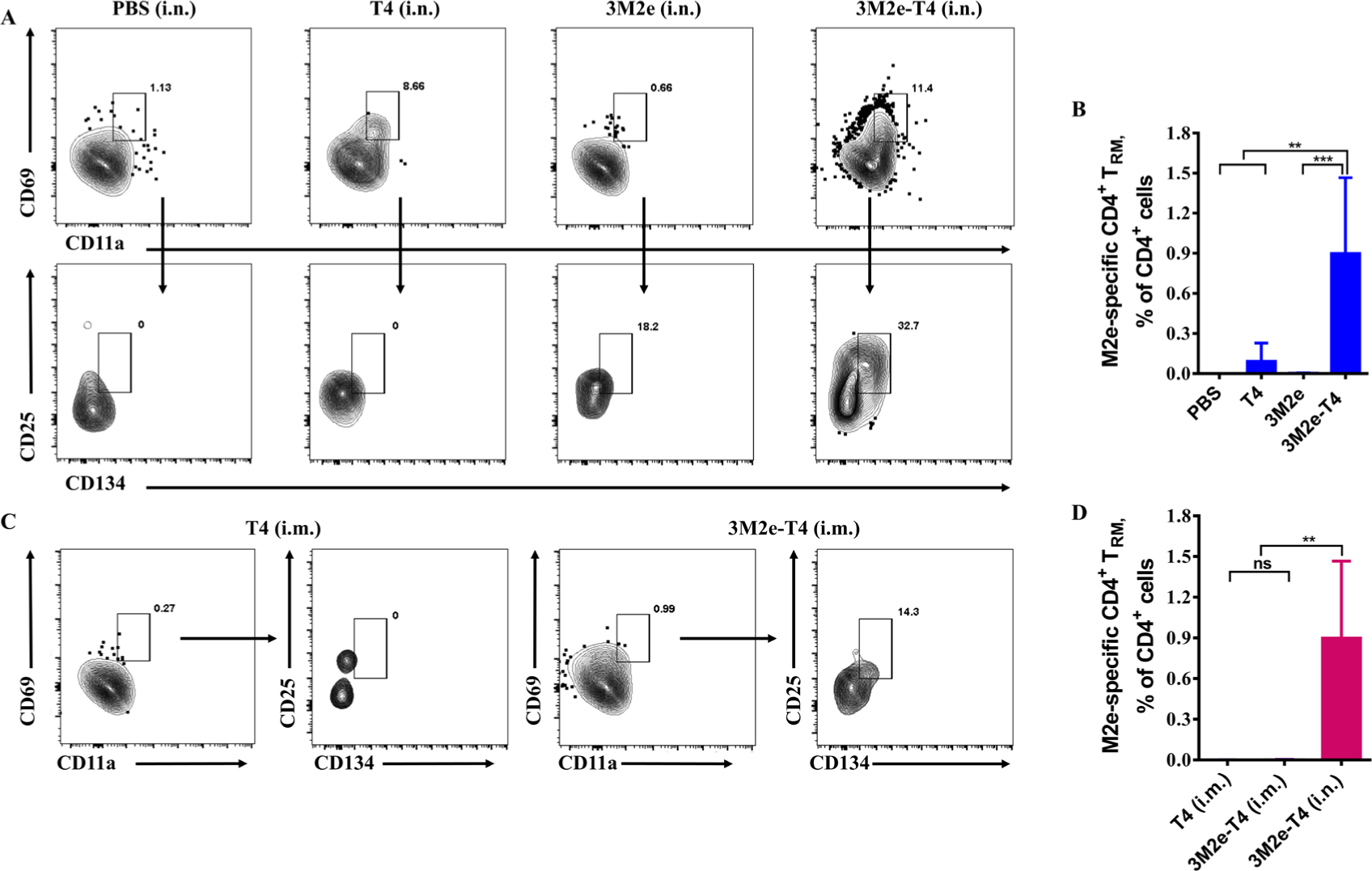
I.n. immunization of 3M2e-T4 nanoparticles elicited M2e-specific CD4^+^ T_RM_ in lungs. CD4^+^ T_RM_ cells in lung were measured at 10 days after administration via i.n. **(A-B)** or i.m. **(C-D)** route. The representative flow cytometry plots **(A, D)** and the frequencies (%) of CD4^+^ T_RM_ in lungs were shown. Data were represented as mean ± S.D. ** P<0.01; *** P< 0.001.

### 2.6 T4 nanoparticle mediated antigen persistence in lungs promoted colocalization of APCs

Previous studies indicated that retention of antigens at the vaccination sites is required for T-cell responses and affinity maturation of antibodies.^[42–44]^ Antigen persistence on mucosal surfaces is also critical for uptake by APCs and subsequent mucosal immune responses, particularly the generation of T_RM_ cells in the lungs.^[12, 45, 46]^ To monitor the persistence of antigens intranasally delivered by T4 phages in mouse, T4 nanoparticles were first labeled with biotin followed by incubation with Alexa Fluor 647-conjugated streptavidin (T4_647_). Strong fluorescence of T4_647_ nanoparticles was observed *in vitro* on an IVIS spectrum imaging system (Figure S3 A-D). After i.n. administrations, fluorescence signals were observed immediately in the noses of mice received T4_647_ nanoparticles but disappeared 40 min later, and no signal was found at any time in the mice administrated with unlabeled T4 phages (Figure 8A). To determine whether T4 nanoparticles enter and persist in the lower respiratory tract, three mice in each group were sacrificed, and the lungs were collected at different time points after i.n. administrations. Fluorescence imaging showed that the signals can be observed as early as 1.5 hr and gradually dropped beyond the detection limit on day 46 after immunization (Figure 8B). Quantification of fluorescent radiant efficiency of each lung also indicated the same trend (Figure 8C). The spleens were also collected from each mouse; however, no fluorescence signals were observed at any time points (Figure S4 A, B). These results indicated that i.n. immunization with T4 nanoparticles leads to antigen deposition and persistence in the lungs with minimal systemic distribution.

**Figure 8.**
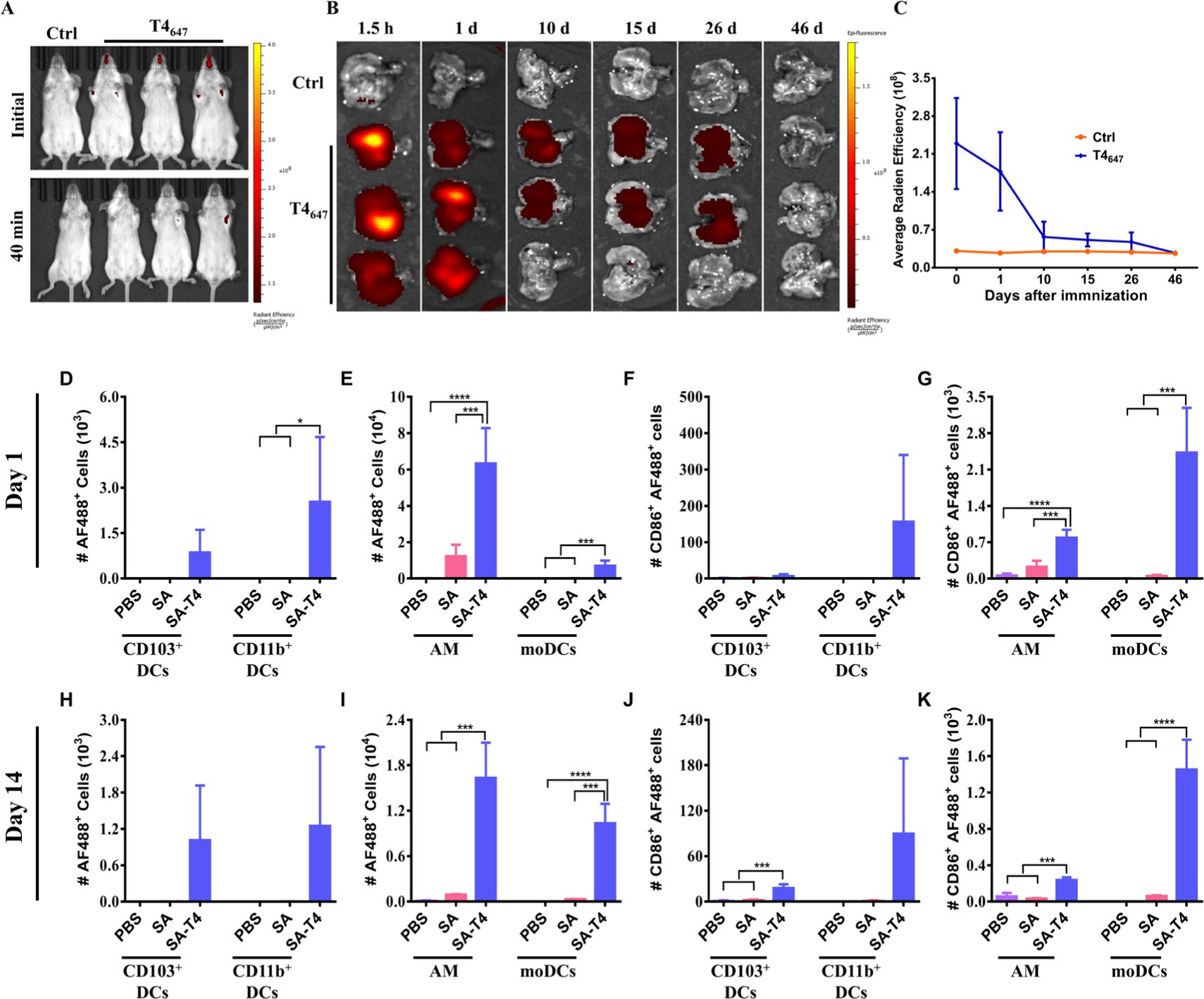
T4-nanoparticles mediated antigen persistence in the lungs and promoted colocalization of antigens and APCs. **(A)** *In vivo* imaging of mice 40 min after i.n. immunization with fluorescently labeled and unlabeled T4 particles as described in Materials and Methods. Lungs were harvested at 1.5 h, 1 d, 10 d, 15 d, 26 d, and 46 d after immunization. Fluorescence imaging **(B)** and quantification of fluorescent radiant efficiency **(C)** of each lung were done using the IVIS spectrum instrument. **(D-K)** Antigens presentation by lung APCs at 1 day, 14 days after i.n. administration of PBS, streptavidin_488_ (SA), and streptavidin_488_-T4 (SA-T4). The number of total streptavidin_488_-positive CD11b^+^ cDCs, CD103^+^ cDCs, AM, and moDCs in lung at 1 day **(D, E)** and 14 days **(H, I)** after immunization was determined using flow cytometer. The total number of CD86^+^ streptavidin_488_-positive CD11b^+^ DCs, CD103^+^ DCs, AM, and moDCs in lung at 1 day **(F, G)** and 14 days **(J, K)** after immunization were also determined. Data were represented as mean ± S.D. * P < 0.05; *** P < 0.001; **** P < 0.0001.

Previous studies indicated that the retention of antigens in lungs leads to the prolonged exposure of the antigen to immune system and enhance the uptake and presentation by immune cells.^[46, 47]^ The lung APCs that initiate adaptive immune responses mainly include CD103^+^ conventional DCs (cDCs) (F4/80^lo^CD103^+^CD11b^lo^), CD11b^+^ cDCs (F4/80^-^CD103^lo^CD11b^hi^), alveolar macrophages (AMs) (CD11b^lo^ F4/80^hi^), and monocyte-derived DCs (moDCs) (CD11b^hi^ F4/80^hi^).^[48–51]^ We asked whether the prolonged exposure of T4 nanoparticles at mucosal surface promotes antigen capture and which kinds of APCs are responsible for antigen capture. To address these questions, mice were immunized with streptavidin_488_-T4, soluble streptavidin_488_, and PBS individually. The Alexa Fluor 488 was used instead of Alexa Fluor 647 to avoid crossover interference between fluorescence-conjugated antibodies and antigens.

On days 1 and 14 after administration, the pulmonary APC subsets were analyzed by flow cytometry after staining with a panel of fluorescence-labelled antibodies against different APC markers (Figure S5). One day after administration, more streptavidin_488_-positive lung CD11c^+^ cells were observed in mice immunized with streptavidin_488_-T4 compared to mice received soluble streptavidin_488_, suggesting that T4 nanoparticles enhance cellular uptake (Figure. S6). Most importantly, significantly higher number of streptavidin_488_-positive CD11b^+^ cDCs, AM, and moDCs were found in mice immunized with streptavidin_488_-T4 than soluble streptavidin_488_ (Figure 8D, E). The same trend was also observed for streptavidin_488_-positive CD103^+^ cDCs although it is not statistically significant (Figure 8D). Both streptavidin_488_-T4 and soluble streptavidin_488_ were mainly captured by AMs (Figure 8E), which is a major APC subset in the lung.^[47, 49, 52]^ CD86 is an activation marker for murine APCs.^[12, 49, 53]^ We found that the streptavidin_488_-T4 nanoparticles induced more CD86^+^ AM and CD86^+^ moDCs one day after vaccination (Figure 8F, G). Similarly, on day 14 after immunization, more streptavidin_488_-positive moDCs and AM were found in mice immunized with streptavidin_488_-T4 compared to the soluble streptavidin_488_ (Figure 8I). Although it is not statistically significant, the same trend was also observed for streptavidin_488_-positive CD103^+^ cDCs and CD11b^+^ cDCs (Figure 8H). Similar results were also observed for streptavidin_488_-positive CD86^+^AM, CD86^+^moDCs, and CD86^+^CD103^+^ DCs (Figure 8J, K).

Antigen-loaded pulmonary APCs will migrate to draining lymph nodes to initiate adaptive immune responses.^[54, 55]^ Meanwhile, antigens passively drained into lymph nodes will be captured by local APCs to activate adaptive immune responses. We, therefore, determined the APCs in mediastinal lymph nodes (MLNs) by accounting the CD11c^+^MHC-II^+^ Streptavidin_488_^+^ cells as described previously.^[49]^ Mice were immunized with PBS, streptavidin_488_-T4 or soluble streptavidin_488_ on day 0, and MLNs were collected on day 1 and 14 to analyze APCs using the gating strategy shown in Figure S7. One day after i.n. administration, the trend towards increased antigen presentation by DCs was found in mice immunized with streptavidin_488_-T4 (Figure 9A). Strikingly, 14 days after immunization, significantly higher number of streptavidin_488_-positive total DCs were observed in streptavidin_488_-T4 group compared to the control group (Figure 9B). We then analyzed streptavidin_488_-positive cDC populations, CD103^+^ cDCs and CD11b^+^ cDCs, because of their critical roles in presenting antigens to T cells in the MLNs.^[54, 56]^ One day after i.n. administration, there is no significant difference between streptavidin_488_-T4 group and soluble streptavidin_488_ group (Fig 9C). The same trend was also found for the activated streptavidin_488_-positive cDC populations (Figure 9D). Strikingly, 14 days after administration, significant higher number of CD11b^+^ cDCs in the MLNs was detected in mice immunized with streptavidin_488_-T4 group compared to the soluble streptavidin_488_ (Figure 9E). The same trend was also observed for streptavidin_488_-positive CD103^+^ cDCs although it is not statistically significant (Figure 9E). Similar results were found for the activated streptavidin_488_-positive CD103^+^ cDCs and CD11b^+^ cDCs (Figure 9F). Interestingly, one day after i.n. immunization, we found that streptavidin_488_-positive cDCs from MLNs in streptavidin_488_ group were mainly CD103^+^ cDCs, while similar amount of CD103^+^ cDCs and CD11b^+^ cDCs were observed in streptavidin_488_-T4 group (Figure S7 A, B). However, 14 days after administration, the CD103^+^ cDCs and CD11b^+^ cDCs were equally activated in both groups (Figure S7 C, D).

**Figure 9.**
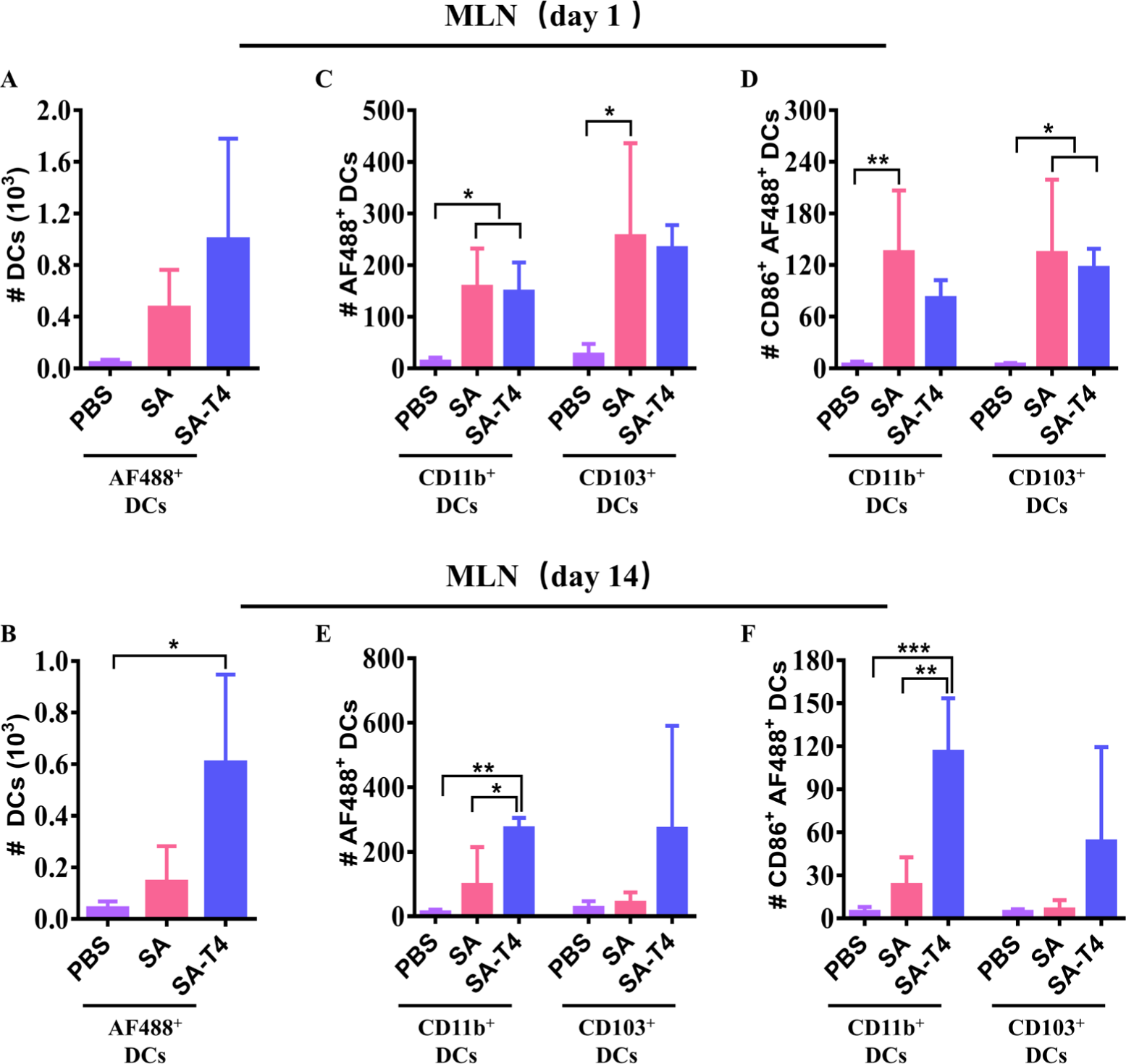
Antigen presentation by DCs in MLNs after i.n. administration of T4-VLP. MLNs were harvested at 1 day and 14 days after i.n. administration with PBS, streptavidin_488_ (SA), and streptavidin_488_-T4 (SA-T4) respectively. The number of total streptavidin_488_-positive DCs, CD11b^+^ cDCs, and CD103^+^ cDCs at 1 day **(A, C)** and 14 days **(B, E)** after immunization was determined using flow cytometer. The number of activated streptavidin_488_-positive CD11b^+^ DCs and CD103^+^ DCs in MLN at 1 day **(D)** and 14 days **(F)** after immunization was also determined. Data were represented as mean ± S.D. * P < 0.05; ** P < 0.01; *** P < 0.001.

Altogether, these results indicated T4-nanoparticles sustained on mucosal surface after i.n. administration, which leads to the prolonged exposure of the antigens to APCs and the enhanced antigen capture. Consequently, more pulmonary DCs were activated and migrated into the MLNs to promote local mucosal immune responses.

## Discussion

Local mucosal immune responses induced by mucosal vaccines combat the invading pathogens at the entry sites and are considered to be the most effective way to prevent infection.^[2]^ A universal vaccine platform that can be customized for the development of mucosal vaccines against any given pathogen is highly desired to prevent infections and transmissions. Here, we demonstrate that the T4 phage nanoparticle could be a promising platform for the development of nasal mucosal vaccines through genetically engineering its genome to generate an antigen-decorated nanoparticle. As a proof of concept, we showed i.n. immunization with 3M2e-decorated T4 nanoparticles efficiently induced local mucosal as well as systemic immune responses and provided mice complete protections against divergent influenza viruses.

Our *in vivo* T4 nanoparticle assembly platform provides a rapid and powerful approach to develop mucosal vaccines against a variety of respiratory pathogens. Compared to *in vitro* assembly, the *in vivo* platform avoids the purification of Soc-antigen fusion proteins and the subsequent *in vitro* assembly. Instead, the antigen coding sequence is inserted into T4 genome in-frame at the C terminus of Soc gene. The recombinant phage can then be generated in ∼2 days using the CRISPR-Cas phage editing technology we recently developed.^[21, 25]^ During the propagation of recombinant T4 phages in *E. coli*, the Soc-antigen proteins will be expressed under the control of native promoter of T4 and self-assembled on T4 capsids to form antigen-decorated nanoparticles.^[21]^ Thus, it is possible to manufacture nanoparticle vaccines in large scale using *E. coli* in a short time at a relatively low cost, which is critical for rapid production of vaccines against the emerging infectious diseases.

The intrinsic adjuvant activity and mucoadhesive property make T4 phage an ideal mucosal vaccine platform. Indeed, we found i.m. immunization with 3M2e-T4 nanoparticles induced robust systemic immune responses without any exogenous adjuvant, which have been extensively confirmed by a series of previous studies.^[15–17]^ Most significantly, we found that i.n. administration with 3M2e-T4 nanoparticles efficiently elicited lung mucosal immune responses in addition to systemic immune responses (Figure 4-7), indicating the mucosal adjuvant activity of T4 phage. This is critical for the development of nasal mucosal vaccines because no adjuvant was approved so far for nasal mucosal use.^[7]^ Furthermore, i.n. delivery of 3M2e-T4 nanoparticles induced enhanced immune responses compared to i.m. immunization and provided mice complete protections against both homologous and heterologous influenza viruses (Figure 3). Further analysis demonstrated that the enhanced protections were because of the lung mucosal immune responses, which were efficiently induced when 3M2e-T4 nanoparticles were administrated via i.n. rather than i.m. route (Figure 5-7). This is consistent with previous finding that mucosal immune responses were efficiently induced only when vaccines were administrated via mucosal routes.^[57]^

One of the biggest challenges for the development of intranasal vaccines is that the delivered antigens must overcome the mucociliary clearance before they were recognized by APCs to initiate local adaptive immune responses. Interestingly, recent studies revealed that T4 phage has mucoadhesive property,^[20]^ which we think might benefit the enrichment of the antigen-decorated T4 nanoparticles on mucosal surface, therefore, increase their chance to be recognized by APCs. In line with this, we found that T4 phage nanoparticles were retained in the lungs for at least 26 days after i.n. delivery (Figure 8B, C). More antigen-positive lung CD11c^+^ cells were observed in the lungs of mice immunized with T4 nanoparticles compared to the soluble antigen even on day one after administration, indicating the enhanced cellular uptake of APCs including CD11b^+^ cDCs, MoDCs, and alveolar macrophages (Figure S6, Figure 8D-E). This trend was also observed on day 14 after immunization (Figure 8H-I), indicating both the nature of particle and the long-term mucosal residency of T4 nanoparticle facilitate antigen uptake by lung local APCs. Antigens that are less than 200 nm can drain to lymph nodes by directly crossing lymphatic vessel walls or be recognized by local APCs and then transported to the draining lymph nodes.^[47, 58]^ Analysis of antigen-positive cDCs in the MLNs, which are the main lymph nodes draining the lung,^[59]^ also revealed the enhanced antigen uptake in T4 nanoparticles (∼80×200 nm) group 14 days after immunization compared to soluble antigens (Figure 9B, E, F). However, on day one post administration, the number of antigen-positive cDCs in the MLNs including CD103^+^ cDCs and CD11b^+^ cDCs is comparable between soluble antigen and T4 nanoparticle groups. These results indicated that the soluble antigens might freely drain to MLNs and are quickly taken up by local DCs at the beginning of immunization, while the T4 nanoparticles that persisted in the lung were continuously taken up by lung DCs, which then migrated to MLNs.

The antigen-specific sIgA antibodies and T cells, in particular T_EM_ and T_RM_ cells, in lung mucosa provide a first line of defense by preventing the entry of respiratory pathogens.^[22, 29, 57, 60]^ Compared to other antibody isotypes, sIgA is more stable in the harsh conditions of mucosal surfaces. The antigen-specific T_EM_ and T_RM_ cells mainly localize in nonlymphoid tissues such as lung, therefore, they respond more quickly than T_CM_ cells against the pathogens that enter body through mucosal tissues.^[22, 61]^ We found that both M2e-specific sIgA and CD4^+^ T cells, including T_EM_ and T_RM_ cells, were efficiently induced by 3M2e-T4 nanoparticles when administrated via i.n. but not i.m. route (Figure 5-7). Our results confirmed previous findings that mucosal immunization and antigen persistence on mucosal surfaces are critical for generation of lung T_RM_ cells.^[12, 45, 46]^ Significantly, we found that only i.n. immunization with 3M2e-T4 induced full protection against heterologous influenza virus (Figure 2 and 3), indicating the M2e-specific mucosal immune responses are critical for cross-protection. Although viral vector vaccines can trigger mucosal sIgA^[62]^ and T_EM_/T_RM_ cells^[63]^ when delivered through i.n. route, particle-based delivery systems such as virus-like particles and nanoparticles are preferred because of the safety concerns. However, adjuvants are needed for most of them in order to efficiently trigger mucosal responses.^[62, 63]^ The *in vivo* assembly T4 vaccine platform allows to rapidly generate a nanoparticle with potent ability to induce mucosal sIgA and T_EM_/T_RM_ cells, probably, because of their intrinsic mucoadhesive property and adjuvant activity.

In conclusion, we demonstrated a quick and powerful approach to develop a mucosal vaccine against the given pathogen using T4 phage platform. The antigen coding sequence can be easily inserted into T4 genome in-frame at the C terminus of Soc gene using the CRISPR-Cas phage editing technology. During the propagation of recombinant T4 phages in *E. coli*, the Soc-antigen proteins self-assemble on T4 capsids to form antigen-decorated nanoparticles with intrinsic adjuvant activity and mucoadhesive property. As a-proof-concept, we showed that the 3M2e-T4 nanoparticles induced potent mucosal immune responses and provided mice complete protection against divergent influenza viruses. Potentially, our platform can be customized for any pathogens and will accelerate the development of future mucosal vaccines.

## Materials and Methods

### Ethics statement

This study was conducted in accordance with the Guide for the Care and Use of Laboratory

Animals recommended by the National Institutes of Health. All animal experiments were performed according to the protocols approved by the Animal Protection and Ethics Committee of Huazhong Agricultural University (HZAUMO-2021-0088), Hubei, China.

### Construction of plasmids

Donor plasmid pModB-Soc-gfp-56, which contains phage T4 genes *ModB* and *56* at 5’ and 3’ end of *gfp* respectively,^[21]^ was used as a vector to construct donor plasmids pSoc-huM2e, pSoc-swM2e, pSoc-avM2e, and pSoc-3M2e (Figure S1A). Briefly, the *gfp* gene in pModB-Soc-gfp-56 was replaced by huM2e, swM2e, avM2e, or 3M2e using conventional molecular cloning. The genes *ModB* and *56* in these donor plasmids are used as two homologous arms to mediate homologous recombination between donor plasmid and T4 genome (Figure S1B). The pLbCas12a-ModB plasmid expressing *ModB*-specific spacers was used to express CRISPR-Cas12 complex that specifically cut T4 genome DNA at the 5’ end of gene ModB (Figure S1B).^[21]^ All primers used in this study were listed in Supplementary Table 1.

### Generation of recombinant T4 phages

The recombinant phages T4 were generated as described previously.^[21]^ Briefly, pLbCas12a-ModB was co-transformed into *E. coli* B834 cells with donor plasmid pSoc-huM2e, pSoc-swM2e, pSoc-avM2e, or pSoc-3M2e respectively. About 300 μL of fresh culture of *E. coli* B834 (10^8^ cfu/mL) was incubated with 10^4^-10^5^ pfu of T4*Δsoc* phage for 7 minutes at 37 °C, which then mixed with 5 mL of top agar and poured onto LB agar plate. After overnight incubation at 37 °C, the generated phage plaques were screened by PCR for the recombinant T4 phages with correct insertion, which were further confirmed by sequencing as described previously.^[21]^

### Purification of antigen-decorated T4 nanoparticles

The propagation and purification of recombinant T4 phages were carried out as previously described.^[16, 21, 64]^ Briefly, a single plaque was transferred to 1.5-mL tube containing 1 mL of Pi-Mg buffer (26 mM Na_2_HPO_4_, 22 mM KH_2_PO_4_, 79 mM NaCl, and 1 mM MgSO_4_). About 300 μL of phage-containing Pi-Mg buffer was incubated with 300 μL of *E. coli* (1.5-2×10^8^ cells/mL) culture at 37 °C for 7 min. About 3 mL of top agar was added to the infection mixture, and were then poured onto LB agar plates. After overnight incubation at 37°C, the whole top agar layer containing phages was transferred using glass slide to 2-L flask containing 1 L of *E. coli* P301 culture (1.5-2×10^8^ cells/mL). After 3-4 h incubation at 180 rpm at 37℃, the phage progenies were collected by centrifugation at 25,000 g at 4 ℃ for 45 min. The phage pellet was resuspended in Pi-Mg buffer containing 10 µg/mL DNase I and 500 μL chloroform. After 30 min incubation at 37 ℃, the phages were further purified by CsCl gradient ultracentrifugation. The purified recombinant T4 phages were analyzed by SDS-PAGE and Western blot using anti-M2e antibodies. The size distribution and zeta potential of recombinant or Hoc^-^Soc^-^ T4 phages were determined using Zetasizer Nano ZS (Malvern Panalytical, UK).

### Immunizations and influenza A virus challenges

Six-to eight-week-old female BALB/c mice were purchased from Laboratory Animal Center of Huazhong Agricultural University, Hubei, China. Soc-3M2e proteins were purified as described previously^[16]^ and antigen-decorated T4 nanoparticles were prepared as described above. Mice were intramuscularly or intranasally immunized with 3M2e-T4 nanoparticles (containing 15 μg 3M2e per dose), a mixture of HuM2e-T4, SwM2e-T4, and AvM2e-T4 (5 μg of each per dose), and soluble Soc-3M2e proteins (15 μg per dose). Mice immunized with PBS or Hoc^-^Soc^-^ T4 were used as controls. For homologous influenza virus challenge, mice were anesthetized with isoflurane and intranasally infected with 5 LD_50_ of A/PR/8/34 (H1N1). For heterologous influenza virus challenge, immunized mice were intranasally infected with 3 LD_50_ of A/Shandong/12/2019 (H3N2). Mice were monitored daily for 14 days for body weight and mortality. Animal with 30% and greater loss in body weight were humanely euthanized immediately and recorded as deceased.

### Antibody detection in sera and BALF

Antigen-specific antibodies levels in sera and bronchoalveolar lavage fluid (BALF) were measured using enzyme-linked immunosorbent assay (ELISA) as described previously.^[16]^ Briefly, serum samples (n=6) were collected on day 0, 14, 28, and 38 as showed in Figure 2A. To collect BALF, mice (n=5) were sacrificed at day 10 after third immunization, and lungs were immediately flushed three times with 1 mL PBS. The BALF was centrifugated at 3,500 g for 10 min to remove any debris before ELISA analysis of M2e-specific antibodies. ELISA plates were coated overnight at 4 ℃ with 200 ng/well of M2e peptide pool containing equal amounts of human, swine, and avian influenza virus M2e peptides. After 5 times washes with PBS-T (PBS containing 0.05% Tween 20), plates were blocked with 3% BSA in PBS-T for 1 h at 37 ℃. Sera and BALF were serially diluted with 1% BSA in PBS-T and added to each well. After incubation at 37 °C for 1 h followed by 5 times washes, peroxidase-conjugated secondary antibody (goat anti-mouse IgG, IgG1, IgG2a, or IgA) was added into each well. After incubation at 37 °C for 1 h, the unbound antibodies were removed by 5 times washes with PBS-T. The 3, 3’, 5, 5’-tetramethylbenzedine (TMB) substrate was added to each well to develop color. Reaction was stopped by 2 M H_2_SO_4_ solution, and absorbance was read at 450 nm.

### Indirect Immunofluorescence Assay

MDCK cells were seeded in 96-well plates at about 2.5×10^5^ cells/well in 100 μL of growth medium (DMEM supplemented with 10% fetal bovine serum), and the plate were incubated at 37 ℃ overnight to allow cells to adhere. The cells were then infected with influenza virus A/PR/8/34 or A/Shandong/12/2019 at MOI of 1. After 1 h incubation at 37 ℃, cells were washed three times with PBS to remove unbound viruses, and serum-free culture media containing 1 μg/mL of TPCK-trypsin was added to each well. After 20 h incubation at 37 ℃, cells were washed with PBS, fixed in 10% formalin for 10 min, and permeabilized with 0.1% Triton X-100 in PBS for 20 min. Cells were blocked with 3% BSA in PBS-T (PBS containing 0.05% Tween 20) at 37 ℃ for 1 h followed by incubation with 1:100 diluted mouse sera for 1 h at 37 ℃. After 5 times washes with PBS-T, cells were incubated with Alexa Fluor 488-conjugated goat anti-mouse IgG (1:1000 dilution, Thermo Fisher Scientific) at 37 ℃ for 1 h. Following five washes with PBS-T, cell nuclei were stained with 1 µg/mL DAPI (BD Biosciences) for 5 min. All samples were imaged using an inverted fluorescence microscope (Thermo Fisher Scientific).

### Measurement of antigen-specific T cell responses in spleen and lung

Spleens and lungs were harvested at 10 days post the third immunization. Single-cell suspensions of spleens were prepared as described previously.^[16]^ Lungs were chopped with scissors into small pieces, which were incubated at 37 ℃ for 1.5 h in 1 mL of RPMI1640 medium containing 10% FBS, 1% penicillin/streptomycin, 1 mg/mL collagenase D (Sigma-Aldrich), and 20 U/mL DNAse I (Thermo Fisher Scientific). Lung single-cell suspensions were then prepared by forcing tissues through 70-μm cell strainers and layered on the top of 40% Percoll solution that overlaid on 80% Percoll solution. After centrifugation at 1000 × g for 20 min, lymphocyte layers were collected, and the cells were washed twice with serum-free RPMI1640 medium. The lymphocytes from spleen and lung were resuspended with 1 mL RPMI1640 supplemented with 10% FBS, 1% penicillin/streptomycin. About 1×10^6^ lymphocytes were seeded to each well of 24-well plates and stimulated with pools of M2e peptides at final concentration of 10 µg/mL. After 44-46 hours of incubation at 37 ℃, 5% CO2, lymphocytes were collected from plates, washed twice with serum-free RPMI1640 medium, and resuspended with staining buffer (PBS with 3% FBS). The live cells were identified by trypan blue staining and stained according to the manufacturer’s protocol (Biolegend, USA). Briefly, cells were blocked with anti-CD32/ CD16 (S17011E, Biolegend) in 100 μL staining buffer for 15 min at 4 ℃. The cells were then incubated for 20 min in the dark with the following fluorophore-conjugated antibodies CD3 (17A2; Biolegend), CD4 (GK1.5; Biolegend), CD44 (IM7; Biolegend) CD62L (MEL-14; Biolegend), CD25 (PC61; Biolegend), CD134 (OX-86; Biolegend), CD11a (M17/4; Biolegend), and CD69 (H1.2F3; Biolegend). Following centrifugation at 250 g for 10 min, cells were washed three times with staining buffer, resuspended with staining buffer, and analyzed by flow cytometry.

### *In vivo* imaging for particle tracking

To biotinylate the Hoc^-^Soc^-^T4 phages, about 2.5×10^11^ phage particles were incubated with biotin at room temperature for 30 min at concentration of 1 mM. Following centrifugation at 23,000 g for 45 min, the unbound biotin was removed, and the biotinylated particles were washed twice with PBS. The biotinylated particles were resuspended in PBS, analyzed by SDS-PAGE and western blot using HRP-conjugated streptavidin, and incubated with streptavidin-conjugated Alex Fluor 647 for 2 h at room temperature. The unbound streptavidin was removed by centrifugation for 45 min at 23,000 g, and the fluorescently labeled particles were washed twice with PBS were visualized by an IVIS Spectrum i*n vivo* imaging system. Mice were then immunized intranasally with fluorescently labeled or unlabeled T4 nanoparticles (1.5×10^11^ particles) and monitored immediately and 40 min after administration using an *in vivo* imaging system. Lungs and spleens were harvested and fluorescence intensity of organs was imaged at 1.5 h, 1 d, 10 d, 15 d, 26 d and 46 d after immunization.

### Analysis of uptake and activation of APCs in pulmonary and MLNs

The T4 phage nanoparticles display Alex Fluor 488-conjugated streptavidin were prepared as described above and intranasally inoculated into mice (streptavidin_488_-T4, 5 μg streptavidin per dose). Mice were immunized with soluble Alex Fluor 488-conjugated streptavidin (5 μg streptavidin per dose) or PBS were used as controls. Mice were euthanized at 1 d and 14 d after immunization, and lungs were flushed with 2 mL PBS containing 1 mM EDTA to collect BALFs. Lungs were then collected after removing the circulating blood cells in the lungs by injecting PBS containing 1 mM EDTA into the right ventricle to flush lung tissues until its color became white. The lung single-cell suspensions were prepared as described above. MLNs were also collected and processed into single-cell suspensions as described above. After removing red blood cells with red blood cell lysis buffer (Solarbio, China), MLN cells were collected by centrifugation at 250 g for 10 min and resuspended in 100 μL staining buffer for staining. The leukocytes in lung single-cell suspensions were isolated by leukocyte isolation kit according to the manufacturer’s instructions (TBD science, China). The leukocytes and BALF were then pooled, and CD11c^+^ fractions were isolated using anti-CD11c magnetic beads according to the manufacturer’s instructions. Briefly, lung leukocytes (≤10^7^ cells) were blocked with 10 μL of anti-CD32/ CD16 (S17011E, Biolegend) for 10 min at room temperature. Ten μL of anti-CD11c magnetic beads (Biolegend) were then added to each tube, which was incubated on ice for 15 min. After centrifugation at 300 g for 10 min, cells were resuspended with 2.5 mL Mojosort^TM^ buffer and placed in the magnet for 5 minutes. After three rounds of cell sorting, the bead-labeled CD11c^+^ cells were pelleted by centrifugation at 250 g for 10 min and resuspended in 100 μL staining buffer for staining. MLN cells and lung CD11c^+^ cells were stained for flow cytometry analysis as described above using following fluorophore-conjugated antibodies DAPI (BD Biosciences), CD45 (30-F11, Biolegend), CD11c (N418, Biolegend), CD11c (HL3, BD Biosciences), MHC Ⅱ (M5/114.15.2, BD Biosciences), F4/80 (T45-2342, BD Biosciences), CD103 (2E7, Biolegend), CD11b (M1/70, BD Biosciences), and CD86 (GL-1, Biolegend).

### Histological analysis of lungs

Mice (n=3) were intranasally immunized and challenged as described above. Five days after challenge with 5 LD_50_ of influenza virus A/PR/8/1934 or 3 LD_50_ of A/Shandong/12/2019, mice were euthanized, and lung tissues were harvested and fixed with 10% formalin. After dehydration using a series of alcohol gradients and paraffin embedding, lung tissues were cut into 4 µm-thick sections. The paraffin-embedded tissue sections were deparaffinized in xylene, rehydrated with graded ethanol, stained with hematoxylin-eosin (HE), and observed under an optical microscope (Nikon, Japan).

### Statistical analysis

All data were analyzed using GraphPad Prism software. Multi-group comparisons were evaluated by one-way analysis of variance (ANOVA). In all cases, differences are considered statistically significant when P values are < 0.05.

## Acknowledgements

This work was supported by grants from National Natural Science Foundation of China [Grant No. 31870915 and 32170094] and Fundamental Research Funds for the Central Universities [Program No. 2662022DKYJ003].

