## Supplementary for "Rapid Development of a Mucosal Nanoparticle Flu Vaccine by Genetic Engineering of Bacteriophage T4 using CRISPR-Cas"

^d^ Hongshan Lab, Wuhan, Hubei 430070, China.

^e^ Institute of Animal Husbandry and Veterinary Sciences, Hubei Academy of Agricultural Sciences, Wuhan, Hubei 430070, China.

^f^ Bacteriophage Medical Research Center, Department of Biology, The Catholic University of America, Washington, DC 20064, USA

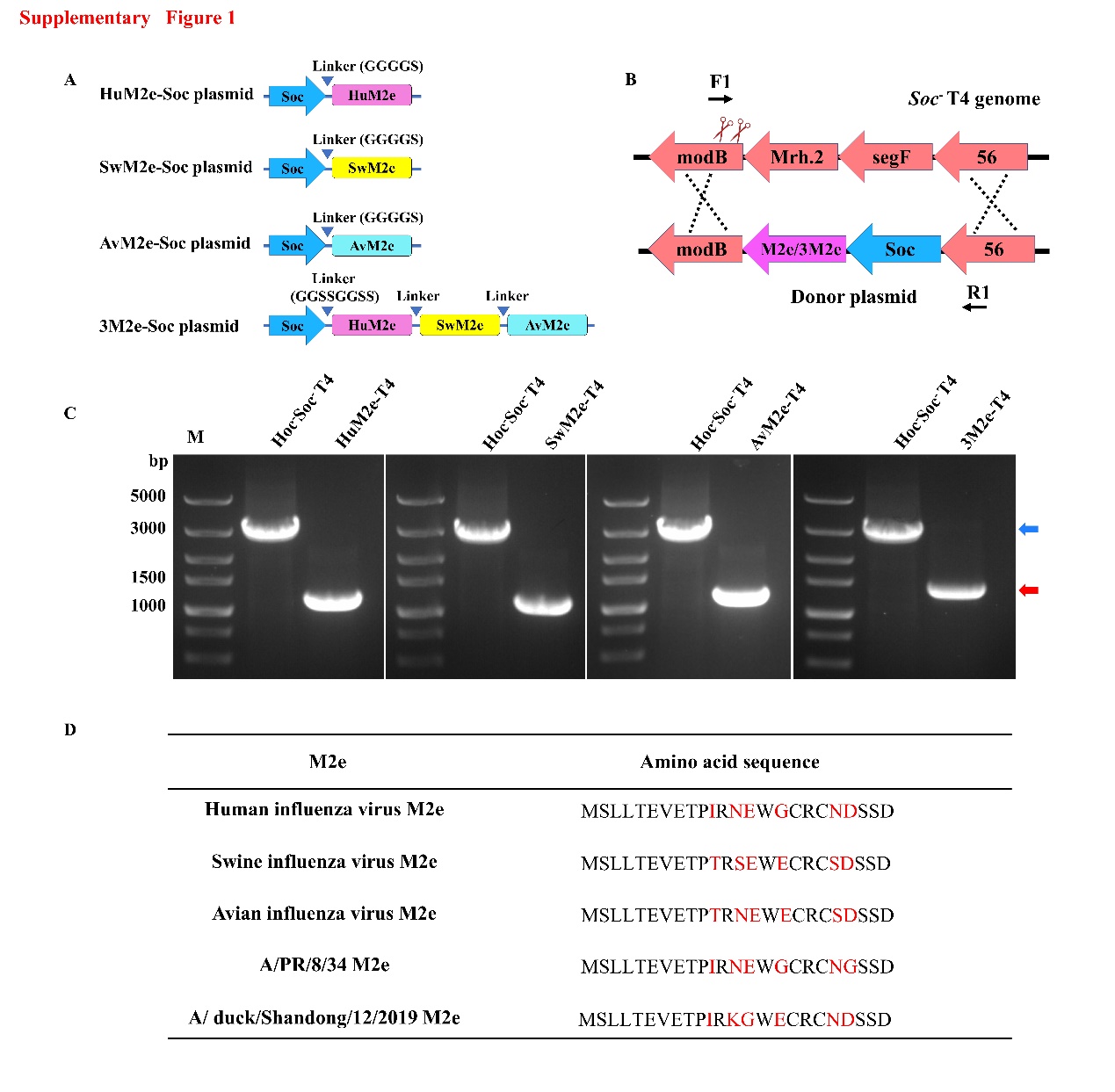
**FIGURE S1**

**FIGURE S1. Analysis of recombinant M2e-T4 mutants. (B)** Schematic showing the fusion M2e gene to the COOH-terminus of the T4 soc gene in T4Δsoc genome. **(C)** The T4 recombinants were analyzed by PCR using primers F1 and R1. Red arrow indicated

HuM2e (1148 bp), SwM2e (1148 bp), AvM2e (1148 bp), and 3M2e (1355 bp). Blue arrow indicated the PCR products of Hoc^-^Soc^-^T4 (3002 bp), which was used as control. **(D)** Amino acid sequences of M2e from human, swine, avian, A/PR/8/34, and A/duck/Shandong/12/2019 influenza virus. Amino acid residues highlighted with red represent differences between different M2e sequences.

**FIGURE S2**

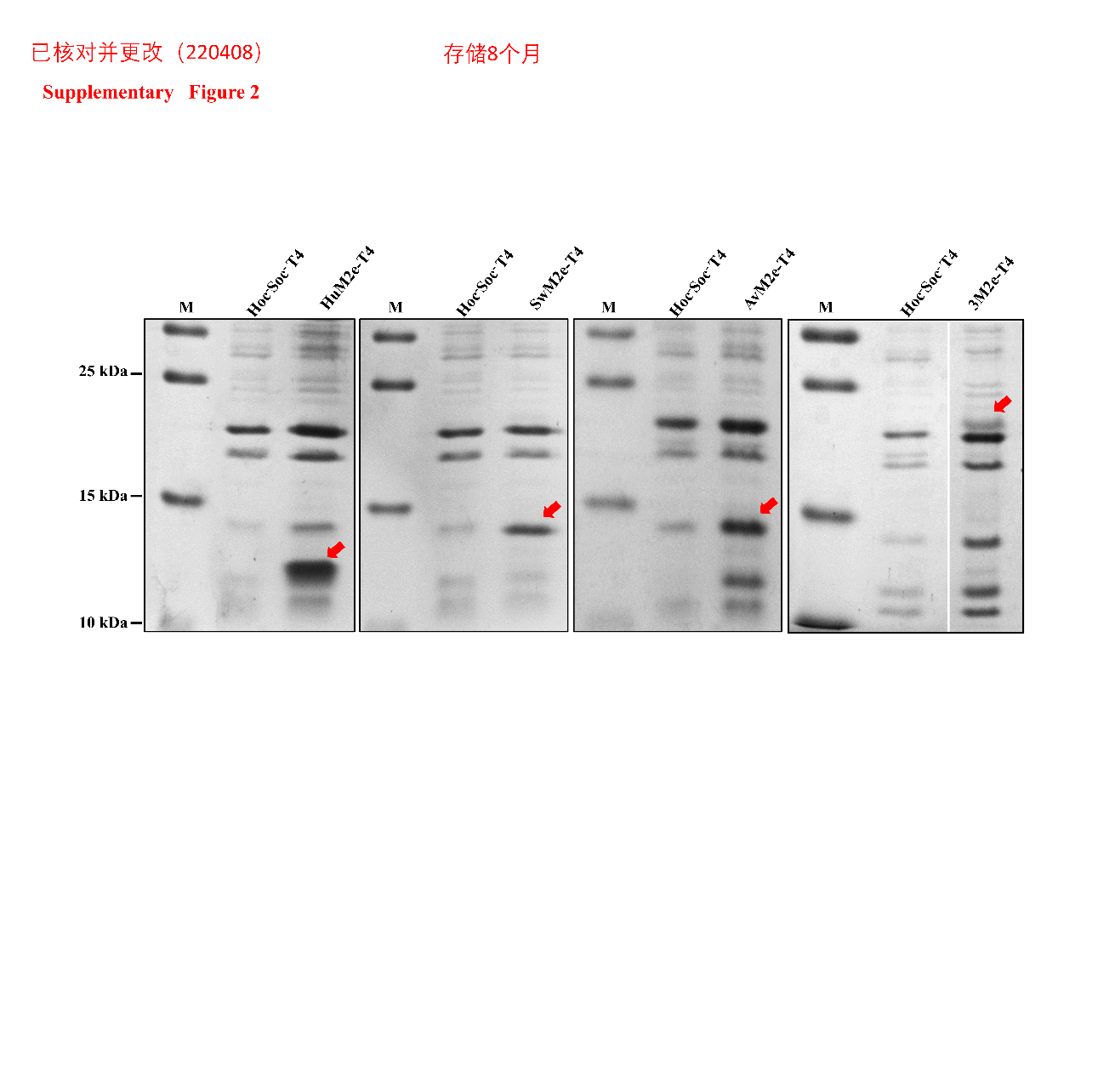

**FIGURE S2. The stability of M2e-T4 nanoparticles.** Purified M2e-T4 nanoparticles were stored at 4 ℃ for 10 months, and analyzed by SDS-PAGE. Hoc^-^Soc^-^T4 was used as control. The bands of Soc-huM2e, Soc-swM2e, Soc-avM2e, and Soc-3M2e proteins were indicated by red arrows.

**FIGURE S3**

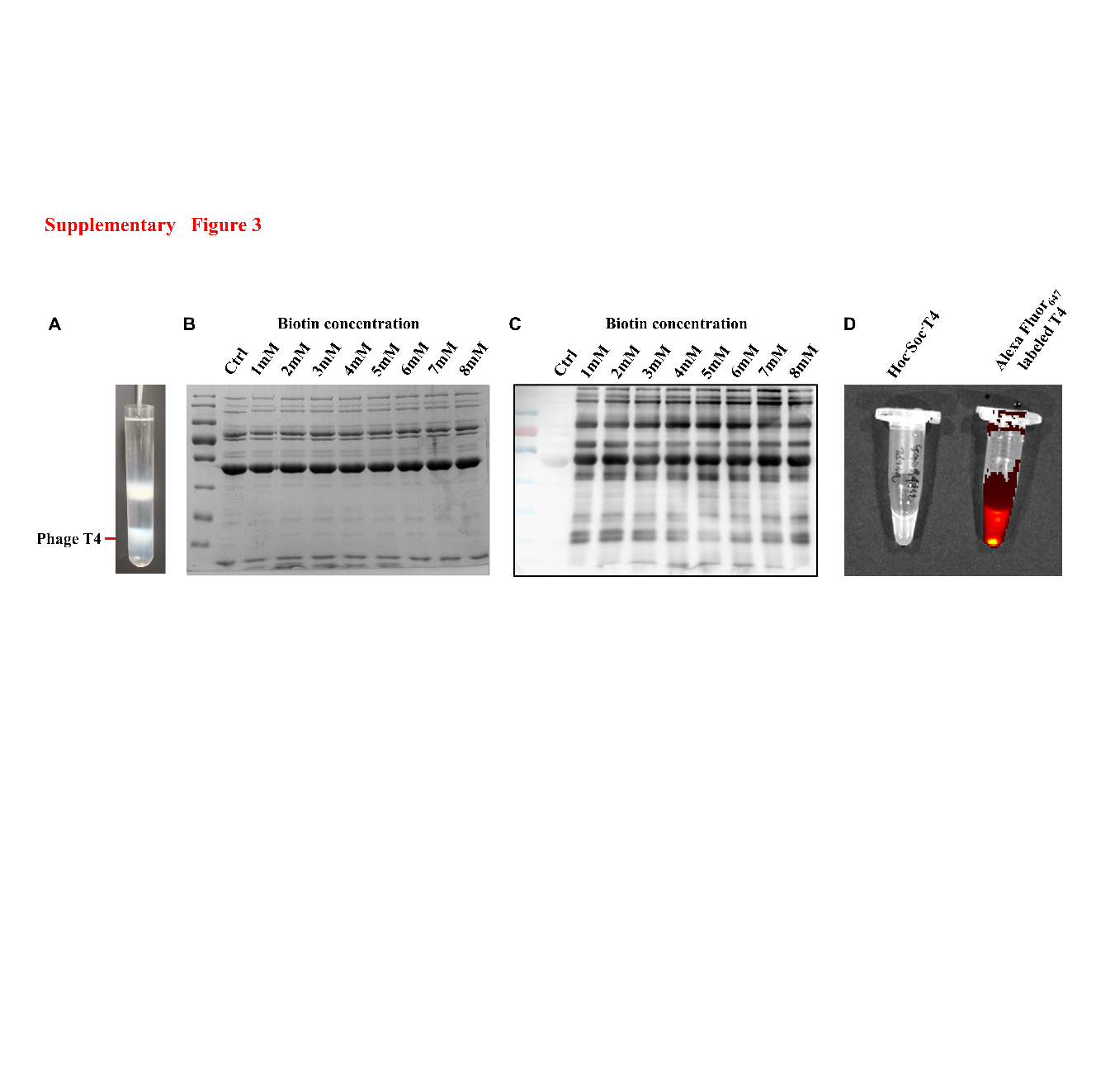

**FIGURE S3. Characterization of fluorescently labeled T4-VLPs. (A)** Hoc^-^Soc^-^T4 phages were purified by CsCl gradient centrifugation. About 2.5×10^11^ Hoc^-^Soc^-^T4 phage were incubated with biotin at increasing concentrations. The biotinylated phages were analyzed by SDS-PAGE **(B)** and streptavidin-HRP western blotting **(C)**. The biotinylated phage were labeled with Alex Fluor 647 conjugated-streptavidin, and fluorescently labeled T4-VLPs were imaged using the IVIS Spectrum instrument **(D)**.

**FIGURE S4**

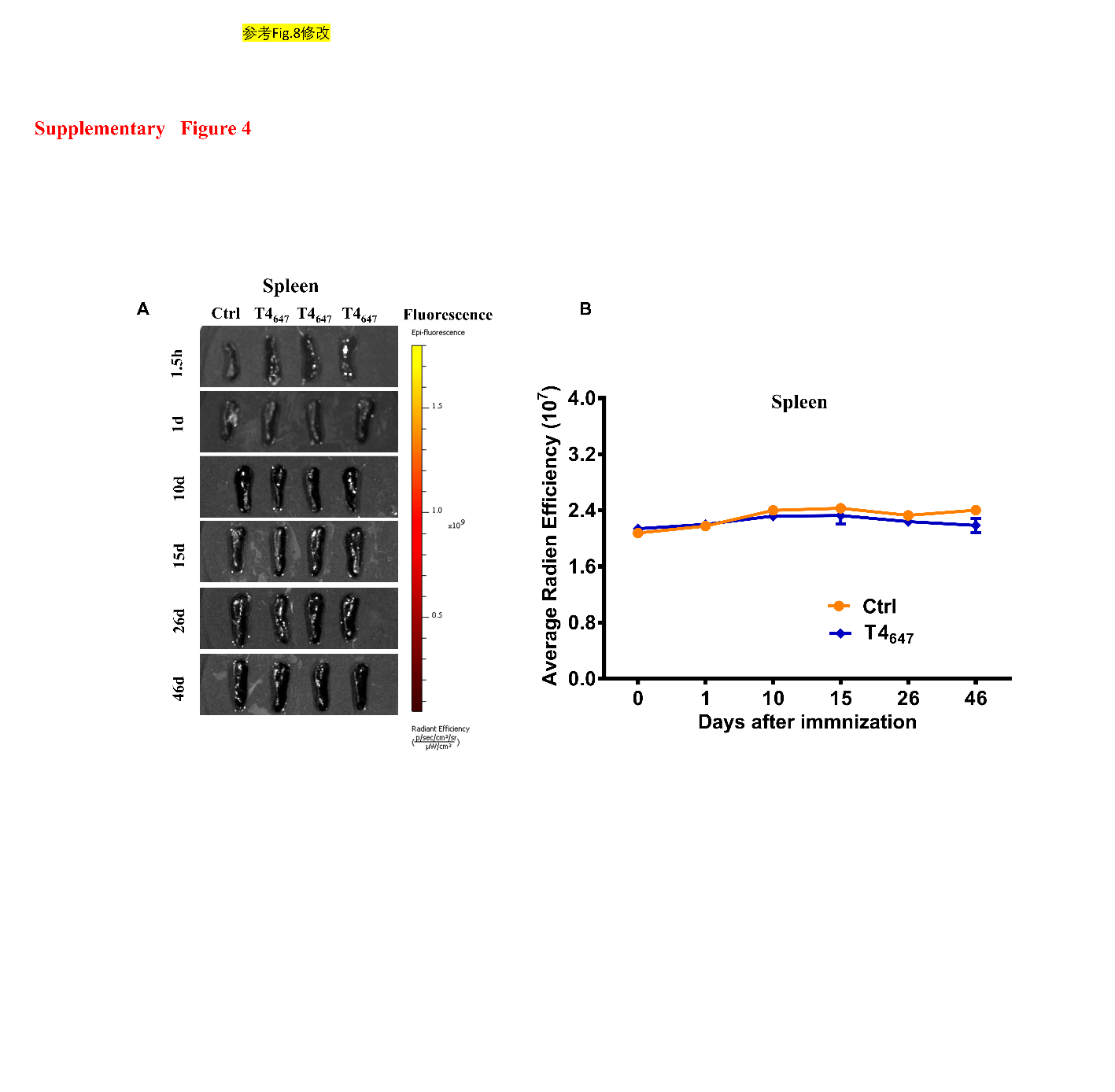

**FIGURE S4. Retention of antigens in spleens.** Spleens were harvested at 1.5 h, 1 d, 10 d, 15 d, 26 d and 46 d after intranasal administration. Fluorescence imaging **(A)** and quantification of fluorescent radiant efficiency **(B)** of each spleen were imaged using the IVIS Spectrum instrument.

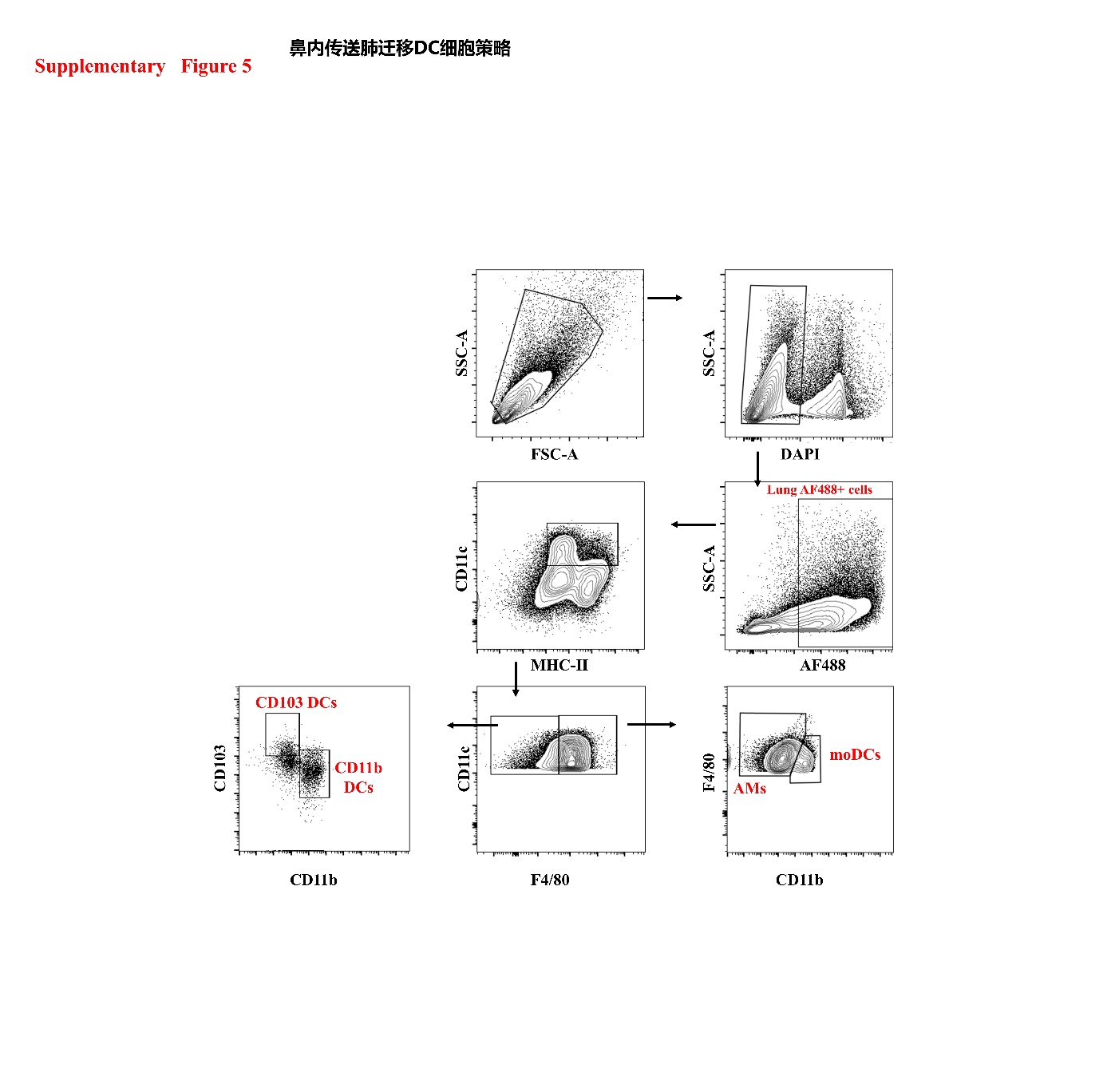
**FIGURE S5**

**FIGURE S5. Gating strategy for the identification of lung APC subsets.**

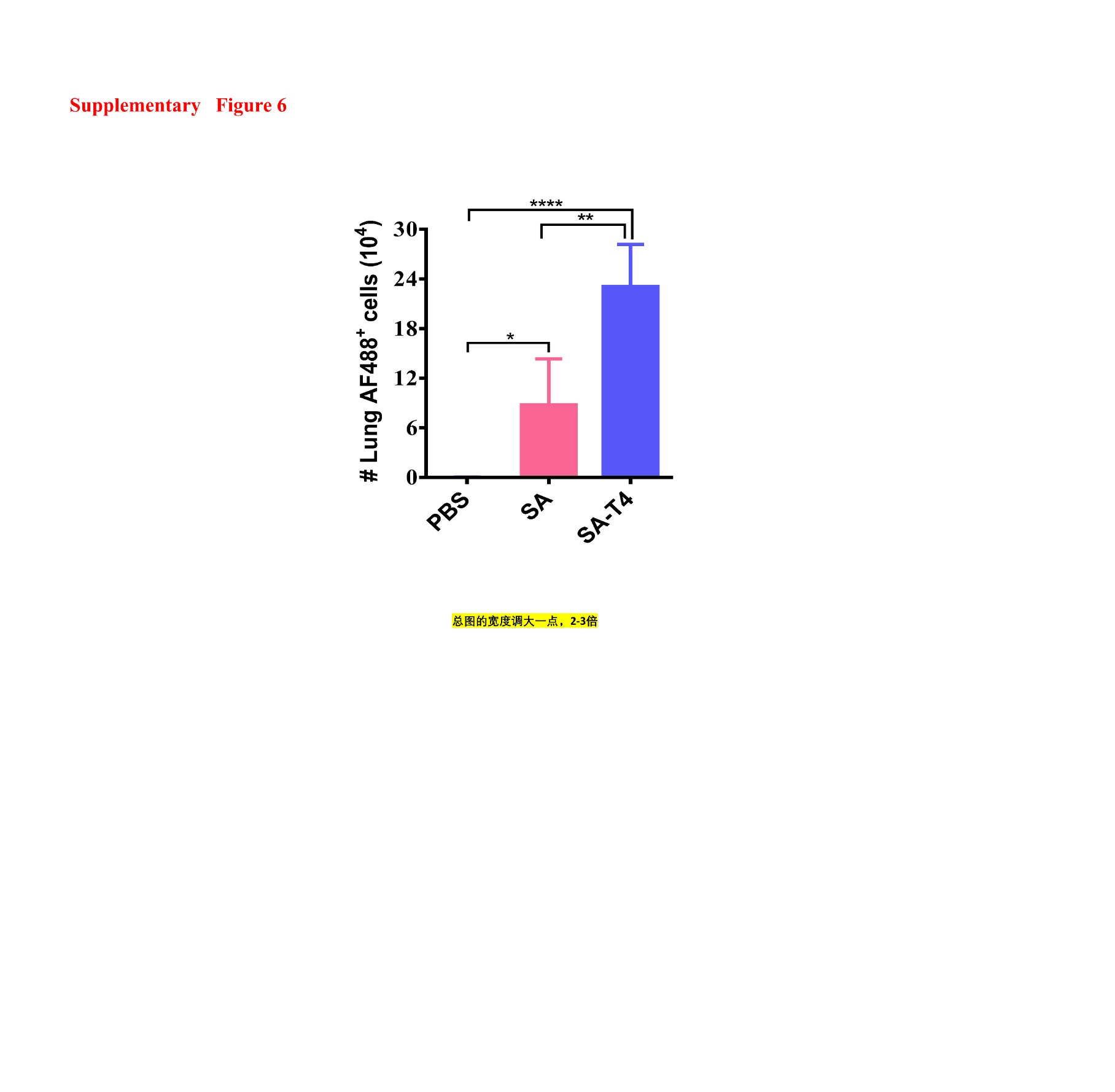
**FIGURE S6**

**FIGURE S6. Analysis of streptavidin^488^-positive cells in lung.** Lungs were harvested at 1 day after i.n. administration with PBS, streptavidin_488_ (SA), and streptavidin_488_-T4 (SA-T4) respectively. The number of total streptavidin^488^-positive CD11c^+^ cells in lung were analyzed.

**FIGURE S7**

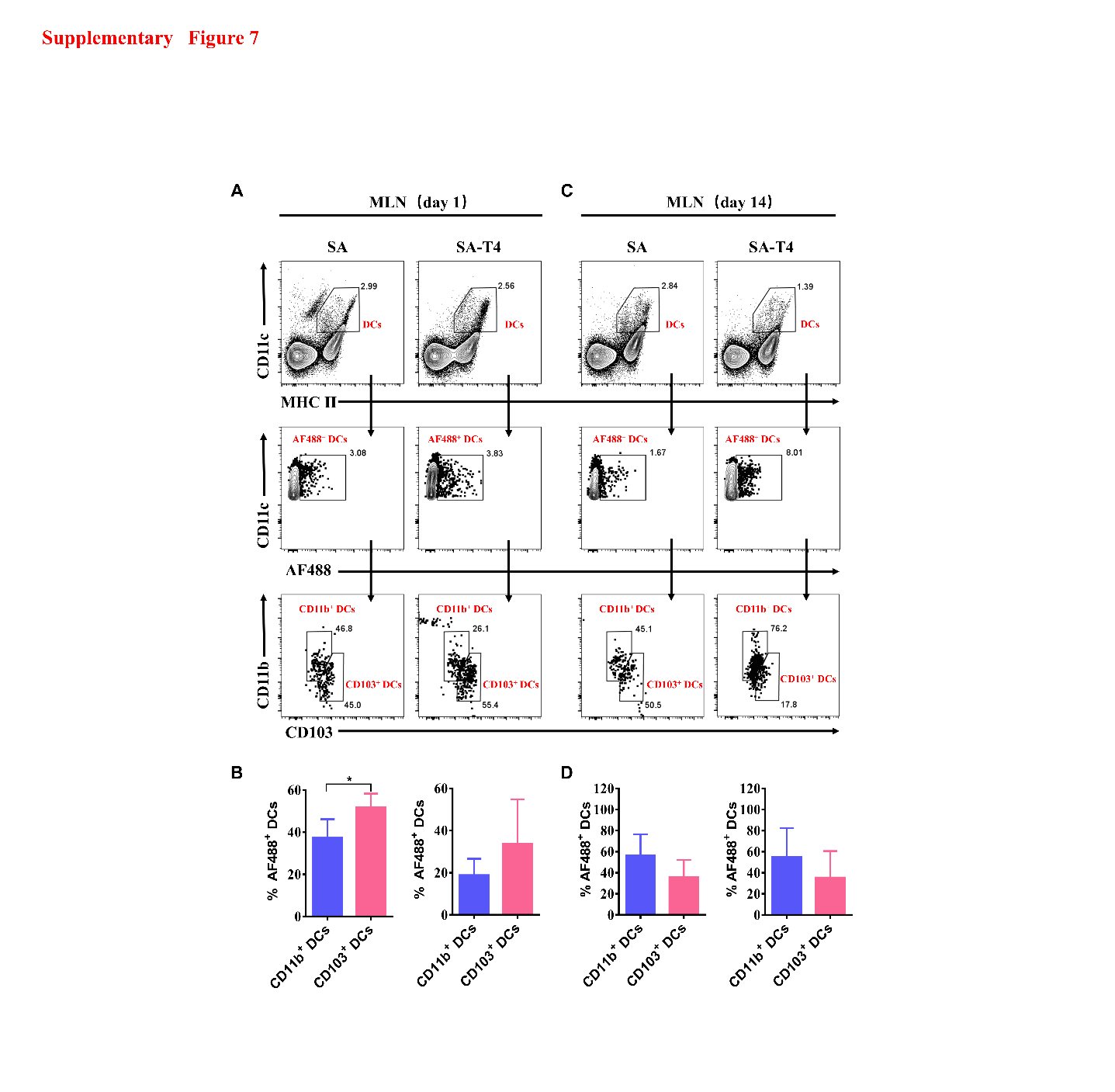

**FIGURE S7. Ratio of antigens captured by DC subtypes in MLNs.** MLNs were harvested and analyzed as described in Materials and Methods. The Representative flow cytometry gating of DC subtypes on day 1 **(A)** and day 14 **(C)** after immunization. Quantification of DC subtype ratio of captured antigen in MLNs on day 1 **(B)** and day 14 **(D)** after immunization.

**Table S1. The sequences of primers used in the experiment**

| 引物 引物序列（5’-3’） | |
| --- | --- |
| R1  F1 | cttggtgaaatgtcacgtggt  gaaagtataatatcgctgatatggt |
| HumanM2e FW1 | AACGAATGGGGCTGCCGTTGCAACGATAGCAGCGATtaataactcaaggactccttcgg |
| humanM2e FW2 | AGCATGAGCCTGCTGACCGAAGTGGAAACCCCGATTCGTAACGAATGGGGCTGCCGTTG |
| HumanM2e FW3 | CCGCTCGAGGGCGGTGGCGGTAGCATGAGCCTGCTGACCG |
| SwineM2e FW1 | AGCGAATGGGAATGCCGTTGCAGCGATAGCAGCGATtaataactcaaggactccttcgg |
| SwineM2e FW2 | AGCATGAGCCTGCTGACCGAAGTGGAAACCCCGACCCGTAGCGAATGGGAATGCCGTTG |
| SwineM2e FW3 | ccgCTCGAGGGCGGTGGCGGTAGCATGAGCCTGCTGACCG |
| AvianM2e FW1 | AACGAATGGGAATGCCGTTGCAGCGATAGCAGCGATtaataactcaaggactccttcgg |
| AvianM2e FW2 | AGCATGAGCCTGCTGACCGAAGTGGAAACCCCGACCCGTAACGAATGGGAATGCCGTTG |
| AvianM2e FW3 | ccgCTCGAGGGCGGTGGCGGTAGCATGAGCCTGCTGACCG |
| modB FW | CCGCTCGAGtaataactcaaggactccttcgg |
